# Global diversity and inferred ecophysiology of microorganisms with the potential for dissimilatory sulfate/sulfite reduction

**DOI:** 10.1101/2023.06.27.546762

**Authors:** Muhe Diao, Stefan Dyksma, Elif Koeksoy, David Kamanda Ngugi, Karthik Anantharaman, Alexander Loy, Michael Pester

## Abstract

Sulfate/sulfite-reducing microorganisms (SRM) are ubiquitous in nature, driving the global sulfur cycle. A hallmark of SRM is the dissimilatory sulfite reductase encoded by the paralogous genes *dsrAB*. Based on analysis of 950 mainly metagenome-derived *dsrAB*-encoding genomes, we redefine the global diversity of microorganisms with the potential for dissimilatory sulfate/sulfite reduction and uncover genetic repertoires that challenge earlier generalizations regarding their mode of energy metabolism. We show: (i) 19 out of 23 bacterial and 2 out of 4 archaeal phyla harbor uncharacterized SRM, (ii) four phyla including the *Desulfobacterota* harbor microorganisms with the genetic potential to switch between sulfate/sulfite reduction and sulfur oxidation, and (iii) the combination as well as presence/absence of different *dsrAB-*types*, dsrL*-types and *dsrD* provides guidance on the inferred direction of dissimilatory sulfur metabolism. We further provide an updated *dsrAB* database including >60% taxonomically resolved, uncultured family-level lineages and recommendations on existing *dsrAB* primers for environmental surveys. Our work summarizes insights into the inferred ecophysiology of newly discovered SRM, puts SRM diversity into context of the major recent changes in bacterial and archaeal taxonomy, and provides an up-to-date framework to study SRM in a global context.

**One sentence summary:** Sulfate/sulfite reducing microorganisms are shaping Earth’s interconnected sulfur and carbon cycles since the Archaean: this legacy unfolds in 27 archaeal and bacterial phyla encountered in diverse marine, terrestrial, and deep-subsurface environments.

## Introduction

The sulfur cycle is one of the most important biogeochemical cycles on Earth (Canfield and Farquhar 2012) tightly interacting with carbon, nitrogen, and metal cycling (Jørgensen 2021). It is mainly regulated by activities of sulfate/sulfite-reducing microorganisms (SRM) and sulfur-oxidizing microorganisms (SOM) as their counterparts (Dopson and Johnson 2012, Wasmund et al. 2017), which cycle sulfur between its most oxidized (sulfate, +IV) and its most reduced state (sulfide, -II). On a global scale, sulfate reduction is one of the dominant processes driving the mineralization of organic matter in anoxic environments. Of the estimated 260 Tmol C_org_ reaching the global seabed each year, one third is mineralized through sulfate reduction in marine sediments (Bowles et al. 2014, Jørgensen 2021). About 90% of the end product, sulfide, is re-oxidized to sulfate either directly or indirectly at the expense of oxygen. This represents 25% of global oxygen consumption in sediments and has a direct impact on the redox state of Earth’s surface. The relevance of sulfur cycling increases further in coastal sediments, where sulfate reduction accounts for 50% of C_org_ mineralization and re-oxidation of sulfide consumes 50% of the oxygen entering the sediment (Jørgensen 2021). While the importance of sulfate reduction in marine environments is well explained by the high availability of sulfate (ca. 28 mM), its role in biogeochemical cycling of anoxic freshwater environments such as lake sediments, groundwater, peatlands, or rice paddy fields is less obvious because of the low prevailing sulfate concentrations (typically 10-300 µM) (Pester et al. 2012). Nevertheless, the rates at which sulfate reduction proceeds can be equally high in marine and freshwater settings, resulting in rapid cycling of sulfur species in anoxic freshwater environments. Because of its less obvious relevance and high variability in space and time, the sulfur cycle in freshwater systems is often referred to as a cryptic or hidden sulfur cycle (Pester et al. 2012). The contribution of sulfate reduction to C_org_ mineralization in anoxic freshwater environments has not been as systematically evaluated as in marine environments, but single studies report values of 17–35% in lake sediments (Urban et al. 1994, Thomsen et al. 2004) and 36–50% in peatlands (reviewed in Pester et al. 2012).

Besides their relevance for biogeochemical cycling, SRM represent an important symbiotic guild in the mammalian intestinal tract (Barton et al. 2017) and are also beneficial in bioremediation, such as degrading hydrocarbons and removing heavy metals from sulfate-containing groundwater and wastewater (Muyzer and Stams 2008, Qian et al. 2019). However, they can also be an economic burden by causing steel corrosion or oil souring (Muyzer and Stams 2008, Rey et al. 2013, Rabus et al. 2015, Singh and Lin 2015, Wolf et al. 2022). In the context of climate change and human-induced eutrophication, it is noteworthy that oxygen concentrations in pelagic zones of the global ocean, coastal waters, and lakes have been declining for decades (Jenny et al. 2016, Breitburg et al. 2018). The resulting oxygen-deficient zones can turn euxinic (anoxic conditions with > 0.1 μM sulfide) upon release of toxic sulfide by SRM, which further aggravates the negative effects of oxygen shortage causing death to fauna including economically relevant fish, shrimp and crabs (Diaz and Rosenberg 2008, Jenny et al. 2016, Bush et al. 2017, Diao et al. 2018, van Vliet et al. 2021).

Most SRM share a canonical core enzyme repertoire for carrying out dissimilatory sulfate reduction (Fig. 1). This intracellular pathway includes the enzymes sulfate adenylyltransferase (Sat), adenylyl phosphosulfate reductase (AprAB), dissimilatory (bi)sulfite reductase (DsrAB), and the sulfide releasing DsrC. The complexes QmoAB(C) and DsrMK(JOP) complement the pathway by transferring reducing equivalents towards AprAB and DsrC, respectively (Pereira et al. 2011, Ramos et al. 2012, Santos et al. 2015). Hereafter, we refer to this pathway as the *dsr*-pathway. Most SRM (with the exception of early diverging archaea) and microorganisms relying on a partial sulfate reduction pathway such as sulfite-, thiosulfate-, and organosulfonate reducers as well as sulfur disproportionating microorganisms utilize in addition DsrD, which is an allosteric activator of DsrAB (Ferreira et al. 2022). Among these enzymes, DsrAB can be used not only as a functional but, with some limitations, also as a phylogenetic marker for SRM. Phylogenetically, this enzyme comprises three major lineages that largely differentiate between (i) reductively-operating DsrAB of archaeal origin, (ii) reductively-operating DsrAB of bacterial origin, and (iii) oxidatively or reverse-operating DsrAB (rDsrAB), which occur in a variety of phototrophic and chemotrophic SOM (Loy et al. 2009, Müller et al. 2015). SOM that rely on rDsrAB for sulfur oxidation also share a number of additional enzymes with SRM, including Sat, AprAB, QmoABC, dsrC, and DsrMKJOP (Dahl 2017, Tanabe and Dahl 2022).

**Figure 1.**
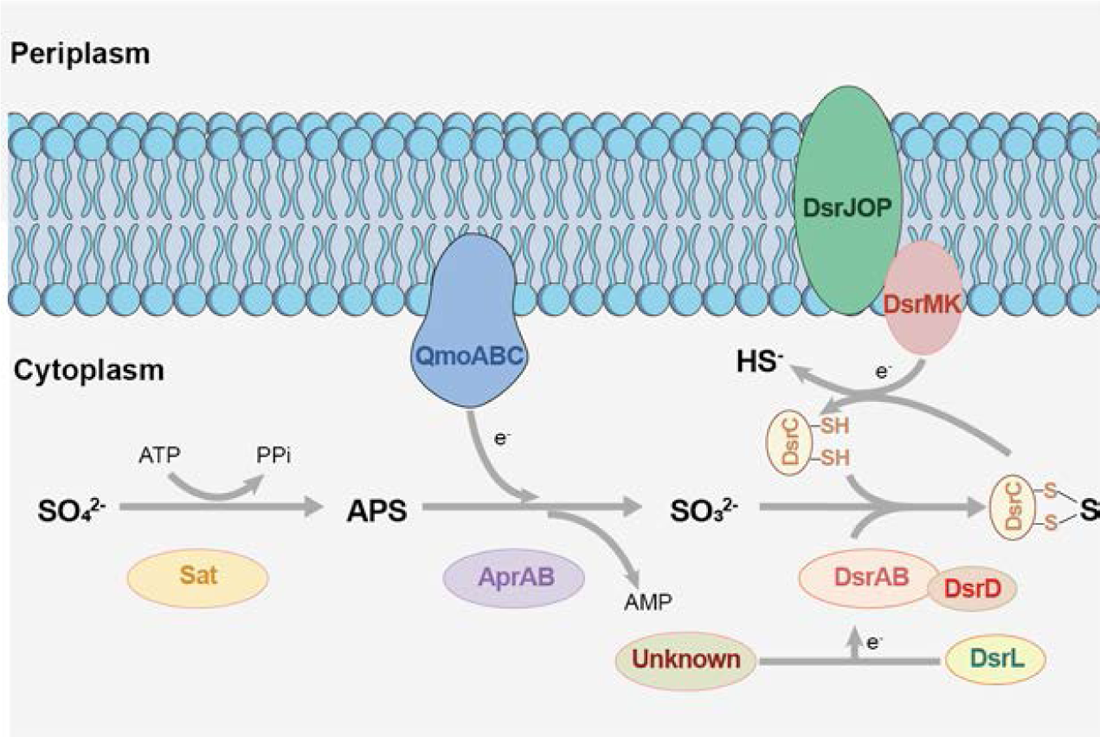
The pathway of dissimilatory sulfate reduction. The *dsr*-pathway includes the enzymes sulfate adenylyltransferase (Sat), adenylyl phosphosulfate reductase (AprAB), dissimilatory sulfite reductase (DsrAB), and the sulfide releasing DsrC protein. The complexes QmoAB(C) and DsrMK(JOP) complement the pathway by transferring reducing equivalents towards AprAB and DsrC, respectively (Pereira et al. 2011, Ramos et al. 2012, Santos et al. 2015). Reducing equivalents required by DsrAB can be delivered by a yet unknown oxidoreductase or DsrL (Löffler et al. 2020). DsrD acts as an allosteric activator of DsrAB in sulfate/sulfite-, thiosulfate-, and organosulfonate reducers as well as sulfur disproportionating microorganisms (Ferreira et al. 2022).

The phylogenetic distinction of reductively and oxidatively operating DsrAB was initially also supported by the presence of additional, presumably SOM-specific enzymes. These include DsrEFH as a sulfur donor protein for DsrC in SOM (Stockdreher et al. 2012) and DsrL as an essential oxidoreductase in sulfur oxidation (Lübbe et al. 2006) that transfers reducing equivalents from rDsrAB to NAD^+^ (Löffler et al. 2020). However, metagenome-assembled genomes (MAGs) from a variety of habitats questioned this clear distinction, with *dsrEFH*, *dsrL*, or both being colocalized together with reductive *dsrAB* (Anantharaman et al. 2018, Hausmann et al. 2018, Tan et al. 2019, Thiel et al. 2019, Ye et al. 2022). The recent identification of two discrete DsrL types, with DsrL-1 occurring only in SOM, while DsrL-2 occurring in organisms with either a reductive/disproportionating or oxidative sulfur metabolism (Löffler et al. 2020), highlights the difficulties in delineating the energy metabolism solely from genomic data. Functional gene prediction is further complicated by the examples of *Desulfurivibrio alkaliphilus* (Thorup et al. 2017) and the so-called cable bacteria affiliated to the *Desulfobulbaceae* (Pfeffer et al. 2012, Risgaard-Petersen et al. 2015). Both can oxidize sulfide by operating the canonical pathway of sulfate reduction in reverse, including a reductive-type DsrAB, and couple this either with intracellular nitrate reduction in case of *D. alkaliphilus* (Thorup et al. 2017) or to electrogenic oxygen or nitrate reduction in spatially separated cells along filaments in the case of cable bacteria (Kjeldsen et al. 2019).

Despite these constraints, *dsrAB*-based molecular approaches have become an important tool for studying the diversity and ecology of SRM in the environment. First introduced by Wagner et al. 1998, cumulative evidence from a large variety of marine, terrestrial, and man-made environments revealed that the diversity of SRM extends massively beyond cultured representatives in the four bacterial phyla *Desulfobacterota* (formerly known as Deltaproteobacteria and Thermodesulfobacteria, Waite et al. 2020), *Bacillota* (formerly known as Firmicutes, Oren and Garrity 2021), *Thermodesulfobiota* (Frolov et al. 2023), and *Nitrospirota* (Oren and Garrity 2021) as well as the two archaeal phyla *Thermoproteota* (formerly known as Crenarchaeota, Oren and Garrity 2021) and *Halobacterota* (formerly part of the Euryarchaeota, Rinke et al. 2021). A systematic review of environmental *dsrAB* genes affiliated with the reductive bacterial-type *dsrAB* revealed at least 13 lineages at the approximate family level that could not be related to any cultured SRM or higher-rank taxa (Pester et al. 2012, Müller et al. 2015). At the species level, a broad census based on *dsrB* amplicon sequencing identified 167,397 species-level operational taxonomic units (OTUs) across 14 different environments (Vigneron et al. 2018). If compared to the approximately 460 described SRM listed in the LPSN database (lpsn.dsmz.de), this means that >99% of SRM diversity is represented by uncultured microorganisms without taxonomic assignment.

Members of well characterized *Desulfobacterota* (*Desulfobacteraceae*, *Syntrophobacteraceae*, *Desulfovibrionaceae*, *Desulfobulbaceae*) often dominate the SRM community in marine and freshwater surface sediments (Vigneron et al. 2018, Wörner and Pester 2019b, Jørgensen 2021) and the uncharted *dsrB* sequence space largely represents low-abundance taxa. However, in certain environments representatives of uncultured *dsrAB* lineages can constitute numerically relevant members of the SRM community (Vigneron et al. 2018), including coastal sediments in the Arctic (Flieder et al. 2021), wetlands (Pester et al. 2012), and deep subsurface marine sediments with active but cryptic sulfur cycling (Leloup et al. 2009), to name a few. Therefore, there is a need to identify these yet unknown SRM and to understand their ecophysiology. In recent years, an increasing number of new *dsrAB*-encoding taxa have been discovered mainly due to metagenomic surveys of environmental samples and the delineation of MAGs. Here, we provide a systematic review of these novel findings, give insights into the increased diversity of (putative) SRM, and place this in the context of the recently proposed overarching changes to bacterial and archaeal taxonomy (Parks et al. 2018, Parks et al. 2020, Rinke et al. 2021, Oren and Garrity 2021). Detailed overviews of well-studied phyla harboring SRM, including cultured and environmental representatives, have been provided in excellent reviews elsewhere (Rabus et al. 2013, Rabus et al. 2015, Langwig et al. 2022).

### Unprecedented diversity of *Bacteria* and *Archaea* with the potential for dissimilatory sulfate/sulfite metabolism

The number of genomes of uncultivated microorganisms assembled from metagenomes is rapidly growing. In recent years, thousands of MAGs from poorly characterized bacterial and archaeal phyla, including those that still lack cultured representatives (candidate phyla), were recovered from a large variety of environments (Anantharaman et al. 2016, Parks et al. 2017, Parks et al. 2018, Rinke et al. 2021). The vast number of novel MAGs allowed researchers to screen for the genomic potential of a dissimilatory sulfur metabolism in microbial lineages that were previously not linked to such processes. In addition, bioinformatics tools were developed lately to identify genes related to sulfur compound dissimilation, transport and intracellular transfer with confidence and in a high throughput manner (Mendler et al. 2019, Neukirchen and Sousa 2021, Tanabe and Dahl 2022, Zhou et al. 2022). This resulted in a burst of discoveries since 2018. For example, a study by Anantharaman et al. (2018) substantially expanded the known diversity of bacterial and archaeal phyla with the capacity for sulfite/sulfate reduction from 7 to 20 phylum-level lineages. Specifically, members of the *Acidobacteriota*, *Armatimonadota*, *Bacteroidota*, *Verrucomicrobiota*, UBA9089 (*Ca*. Desantisbacteria), SAR324 (*Ca*. Lambdaproteobacteria), *Ca*. Zixibacteriota and *Ca*. Hydrothermarchaeota contained *dsr*-pathway genes to perform sulfate/sulfite reduction (Anantharaman et al. 2018). *Chloroflexota* associated with marine sediments (Wasmund et al. 2016) and freshwater subsurface sediments (Hug et al. 2016), newly described members of the *Nitrospirota* recovered from rice paddy soil (Zecchin et al. 2018), enigmatic bacteria of novel candidate phyla such as SZUA-79 (*Ca*. Acidulodesulfobacterales) from an artificial mine drainage with very low pH (pH ∼2) (Tan et al. 2019), and cryptic candidate phyla like UBA9089 and CG2-30-53-67 (Probst et al. 2017) contribute further to the diversity of bacteria with the potential to reduce sulfate/sulfite. Anaerobic oxidation of methane (AOM) or other alkanes coupled to sulfate reduction was suggested to be performed by microbial consortia of methanotrophic archaea and sulfate-reducing bacteria (Boetius et al. 2000, Knittel and Boetius 2009). The recent finding that some *Halobacterota* (*Archaeoglobaceae*) and *Thermoproteota* (*Ca*. Methanodesulfokores washburnensis) encode both a methanogenesis pathway and the capability to perform sulfate or sulfite reduction, respectively, suggests that sulfur-dependent AOM can be carried out in a single organism, independent from syntrophic interactions (McKay et al. 2019, Wang et al. 2019).

The enormous increase in the phylogenetic breadth of bacteria and archaea with the potential for DsrAB-based dissimilatory sulfate/sulfite reduction is currently missing a systematic overview. Here, we screened all publicly available and functionally pre-annotated genomes and MAGs summarized on the Annotree platform v.1.2.0 (Mendler et al. 2019) for the presence of *dsrAB* (http://annotree.uwaterloo.ca, accessed on April 3^rd^, 2023, for bacteria and February 6^th^, 2023, for archaea). This resulted in a total of 902 bacterial and 48 archaeal genomes distributed across 27 and 4 phyla according to the GTDB-Tk taxonomy (Parks et al. 2018, Rinke et al. 2021), respectively (Fig. 2, Fig. 5). Phyla provisionally split by the GTDB release 214 into different sublineages such as Bacillota and Bacillota_A to Bacillota_H were counted as one phylum. Since Annotree (Mendler et al. 2019) is based on the Genome Taxonomy Database, which considered only MAGs of >50% completeness and >10% contamination (Parks et al. 2018, Rinke et al. 2021), MAGs that did not fulfill these quality criteria were omitted from our analysis. These included, for example, a *Verrucomicrobiota* MAG with early diverging *dsrAB* (Verrucomicrobia bacterium SbV1) as well as representatives of the Schekmanbacteria (Schekmanbacteria bacterium RBG_13_48_7) and *Chloroflexota* (Chloroflexi bacteria RBG_13_60_13 and RBG_13_52_14) described previously in the literature (Anantharaman et al. 2018).

**Figure 2.**
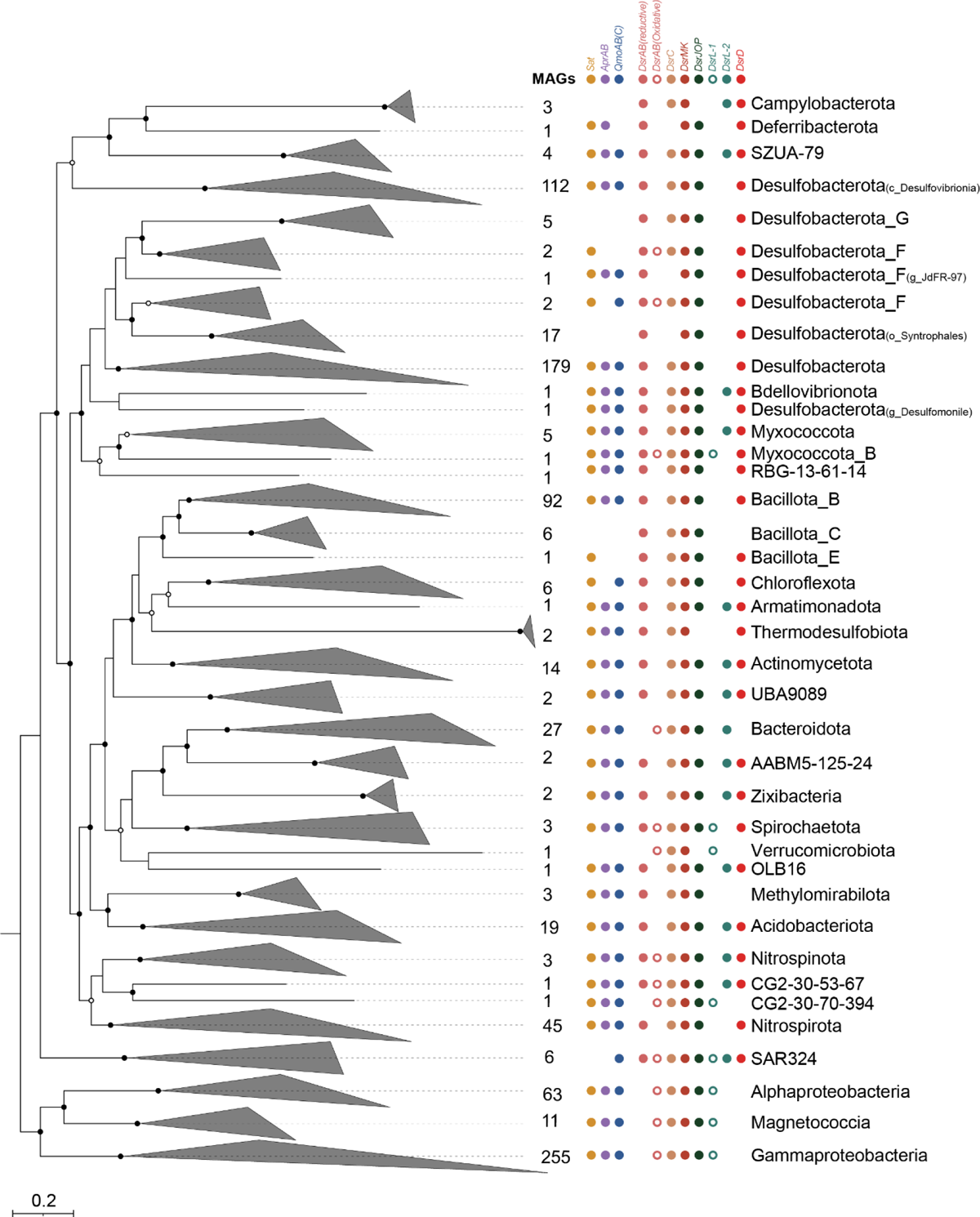
Phylogeny and *dsr*-pathway composition of *dsrAB*-encoding bacteria. Bacterial genome tree inferred from 120 concatenated proteins as based on the GTDB taxonomy (Parks et al., 2018). The phylogenomic tree was inferred from 902 bacterial (metagenome-assembled) genomes. The scale bar indicates 20% sequence divergence. The maximum likelihood tree was constructed with IQ-TREE (Nguyen et al. 2015) using automatic substitution model selection (LG+F+R10) and ultrafast bootstrap analysis (n=1000). Bootstrap support is indicated by black dots (≥90%) or black circles (70–90%). Besides *dsrAB*, the presence of *dsr*-pathway genes was indicated if >30% of *dsrAB*-containing genomes encoded the respective genes as inferred by an automated hmm search (Zhou et al. 2022, custom-made pHMM for DsrL).

The identified *dsrAB*-encoding genomes and MAGs were analyzed in more detail using protein hidden Markov Models (pHMMs) (Zhou et al. 2022, custom-made pHMM for DsrL) to search for genes encoding proteins known to be involved in dissimilatory sulfur metabolism (Fig. 1, Supplementary Tables S1). About half of the identified bacterial genomes were affiliated to the *Desulfobacterota* (35%), *Bacillota* (11%), *Nitrospirota* (5%), and *Thermodesulfobiota* (0.2%), which together represent phyla encompassing all cultured and well-characterized bacterial SRM (Rabus et al. 2013, Rabus et al. 2015). In addition, members of the *Campylobacterota* (i.e. *Desulfurella* spp.) represented cultured thiosulfate-reducing bacteria that employ the *dsr*-pathway (Florentino et al. 2017, Florentino et al. 2019). The second largest group of *dsrAB*-encoding bacterial genomes belonged to the *Pseudomonadota* (36%), including the classes *Alphaproteobacteria, Gammaproteobacteria,* and *Magnetococcia*, and to the *Bacteroidota*, class *Chlorobia* (2%). The two phyla represent cultured and well-characterized SOM with an oxidatively-operating *dsr*-pathway (Loy et al. 2009, Dahl 2017). Members of *dsrAB*-encoding, canonical SRM- or SOM-related lineages also encoded all other *dsr*-pathway genes. The only exception to this was *dsrL*, either of type 1 or 2, which was typically encoded by canonical SOM-related lineages but was absent in the overwhelming majority of genomes affiliated to canonical SRM-related lineages, underlying its relevance for sulfur oxidation (Lübbe et al. 2006). Vice versa, *dsrD* was typically encoded in canonical SRM-related but not in SOM-related lineages, underlying its relevance for sulfate/sulfite reduction (Ferreira et al. 2022) (Fig. 2).

The remaining 11% of *dsrAB*-encoding genomes were spread over 20 different bacterial phyla with the *Acidobacteriota* (19 MAGs) and *Actinomycetota* (14 MAGs) representing the most prominent groups (Fig. 2). Representatives of the *Acidobacteriota, Zixibacteria, Bdellovibrionota, Armatimonadota,* and the candidate phyla UBA9089 (Desantisbacteria), SZUA-79, OLB16, and AABM5-125-24 were all characterized by the full set of *dsr-*pathway genes, including *dsrD* as indicator for a reductively operating metabolism and *dsrL* of type 2 (Fig. 2), which is present in organisms with either a reductive/disproportionating or oxidative sulfur metabolism. The same was true for MAGs within the *Actinomycetota* and *Myxococcota*, with three exceptions which are described in more detail below. Representatives of the *Chloroflexota, Deferribacterota*, and candidate phylum RBG-13-61-14 also encoded *dsrD* but lacked *dsrL*. Since most of the other *dsr-* pathway genes could be recovered for the latter three phyla, a reductively operating pathway is indicated here as well. Members of the *Methylomirabilota* (previously assigned to *Candidatus* Rokubacteria, Anantharaman et al. 2018) lacked both *dsrD* and *dsrL* but belong to the group of microorganisms with the earliest diverging DsrAB (Fig. 2). This group of bacteria and archaea with early diverging DsrAB consistently lacks *dsrD* and *dsrL* but contains cultured representatives with a reductively operating *dsr-*pathway (Anantharaman et al. 2018, Ferreira et al. 2022). In summary, members of fourteen phyla without cultured SRM (representing 7% of all recovered bacterial genomes) encode the full enzyme complement to perform dissimilatory sulfate/sulfite reduction. In contrast, representatives of the *Verrucomicrobiota* and candidate phylum CG2-30-70-394 lacked *dsrD* but encoded *dsrL* of type 1, which resembles the situation in canonical SOM and is indicative of an oxidatively operating sulfur metabolism.

The situation was more complex in members of the *Nitrospirota, Nitrospinota*, *Spirochaetota*, *Bacteroidota,* and SAR324. In these five phyla, different MAGs of the same phylum encoded different gene combinations of the *dsr-*pathway. Within the *Nitrospirota*, the majority of MAGs encoded *dsrD* but lacked *dsrL*, including cultured SRM of the genus *Thermodesulfovibrio* (Zecchin et al. 2018). However, there were two MAGs that lacked *dsrD* but encoded either *dsrL*-1 (f_RBG-16-64-22) or *dsrL*-2 (f_9FT-COMBO-42-15) indicating an oxidatively operating *dsr*-pathway (Fig. 3, Supplementary Table S1). The opposite was true for the *Bacteroidota*. Here, most MAGs and genomes of cultured representatives belonged to the class *Chlorobia* (family *Chlorobiaceae*), which represent canonical SOM and encode gene combinations of the *dsr*-pathway typical for SOM (no *dsrD*, *dsrL* of type 1 or 2). However, five representatives of the *Bacteroidota* family UBA2268 (class *Kapabacteria*), encoded both *dsrD* and *dsrL* of type 2 (which was clearly distinct from *dsrL* of the *Chlorobiaceae*) indicative of a reductively operating *dsr*-pathway. *In situ* transcriptional profiles of UBA2268 MAGs recovered from hot spring microbial mats clearly supported a reductively operating *dsr*-pathway activated under anoxic conditions (for details see below, Thiel et al. 2019). For members of the *Nitrospinota* and *Spirochaetota*, MAGs were more evenly distributed and either encoded gene combinations indicative of a reductive (*dsrD* along with *dsrL*-2 or no *dsrL*) or oxidative sulfur metabolism (no *dsrD* but *dsrL* of type 1 or 2). Furthermore, our analysis recovered six SAR324 members with different gene combinations of the *dsr*-pathway. Two SAR324 MAGs affiliated to the provisional family XYD2_FULL-50-60 were recovered from the terrestrial subsurface with an indicated reductive sulfur metabolism (*dsrD* along with *dsrL*-2). The remaining four MAGs (f_NAC60-12) were recovered from marine environments with an indicated oxidative sulfur metabolism (no *dsrD*, *dsrL*-1). The latter coincides well with reports of *dsr*-pathway encoding SAR324 from oxygenated deep ocean waters (Swan et al. 2011), hydrothermal vent plumes (Sheik et al. 2014), and marine oxygen minimum zones (van Vliet et al. 2023).

**Figure 3.**
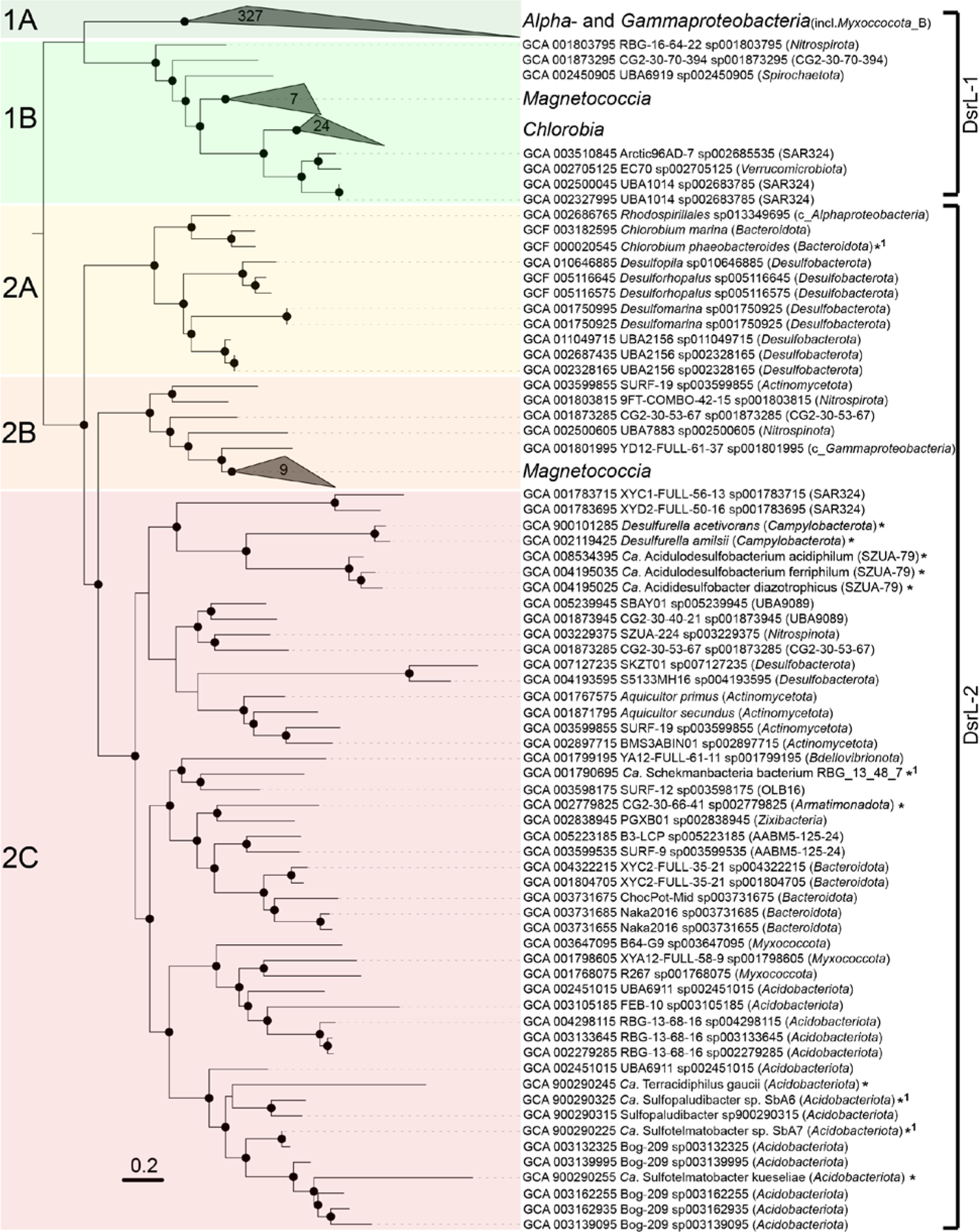
Phylogenetic analysis of bacterial DsrL proteins. The maximum likelihood tree was inferred from 438 DsrL proteins and constructed with IQ-TREE (Nguyen et al. 2015) using automatic substitution model selection (Q.pfam+I+R9) and ultrafast bootstrap analysis (n=1000). Bootstrap support is shown by black dots (≥90%). DsrL sequences marked with an asterisk were not detected by the custom-made pHMM, but have been previously described in the literature (Hausmann et al. 2018, Löffler et al. 2020). Additional DsrL sequences (*1) were collected from MAGs with low completeness, which were not included in our MAG analysis.

An unusual gene combination was encoded by eight MAGs spread over the phyla *Actinomycetota*, *Myxococcota*, CG2-30-53-67 (one MAG each) and the *Desulfobacterota* (five MAGs within the family *Desulfocapsaceae*). Here, at least two sets of bacterial-type *dsrAB,* including always one reductive and one oxidative one, were encoded on the same genome (Fig. 4, Table S1). The most striking example was *Actinomycetota* MAG GCA_003599855, which was recovered from a 1.5 km deep terrestrial aquifer (Momper et al. 2017) and encoded two sets of oxidative *dsrAB* and one set of reductive *dsrAB*. On a single contig, both reductive and oxidative *dsrAB* were encoded in close proximity. The reductive *dsrAB* were encoded upstream of *dsrD* and *dsrL*-2, with the latter falling into the major DsrL-2 cluster. The oxidative *dsrAB* were encoded just four genes further downstream and flanked by *dsrEFH* (upstream) and a second copy of *dsrL*-2 (downstream). The latter clustered together with DsrL-2 of *Magnetococcia, Nitrospinota* family UBA7883 and *Nitrospirota* family 9FT-COMBO-42-15, which all have a verified or indicated oxidative sulfur metabolism (Löffler et al. 2020). A second contig carried the third *dsrAB* set of oxidative type just upstream of a fragmented *dsrL*-2 located at the end of the contig (Fig. 4). From the same terrestrial subsurface habitat, Myxococcota_B MAG GCA_003598065 (Momper et al. 2017) was recovered as another interesting example. It encoded reductive *dsrAB* downstream of *dsrD* and upstream of *dsrCTMKJOP* and on a separate contig oxidative *dsrAB* just upstream of *dsrEFHCMKLLJOP*. Interestingly, *dsrL* was of type 1 and present in two copies in direct vicinity. An additional Myxococcota MAG (GCA_003153055) with oxidative *dsrAB* was recovered as well but was missing further genes indicative of a reductive or oxidative metabolism. Yet another unusual representative of the terrestrial deep subsurface was the single MAG representing candidate phylum CG2-30-53-67 (Table S1), which was recovered from deep groundwater (Probst et al. 2017). Also here the reductive *dsrAB* were encoded upstream of *dsrD* and *dsrL-*2, with the latter falling into the major DsrL-2 cluster, and the oxidative *dsrAB* were encoded upstream of a second copy of *dsrL-2*, which clustered together with *dsrL*-2 of *Magnetococcia* (Fig. 3, Fig. 4). As suggested before (Löffler et al. 2020), this bacterium is likely capable of switching the direction of dissimilatory sulfur metabolism by regulating the different types of DsrABL. The same is likely true for the *Actinomycetota* MAG described above.

**Figure 4.**
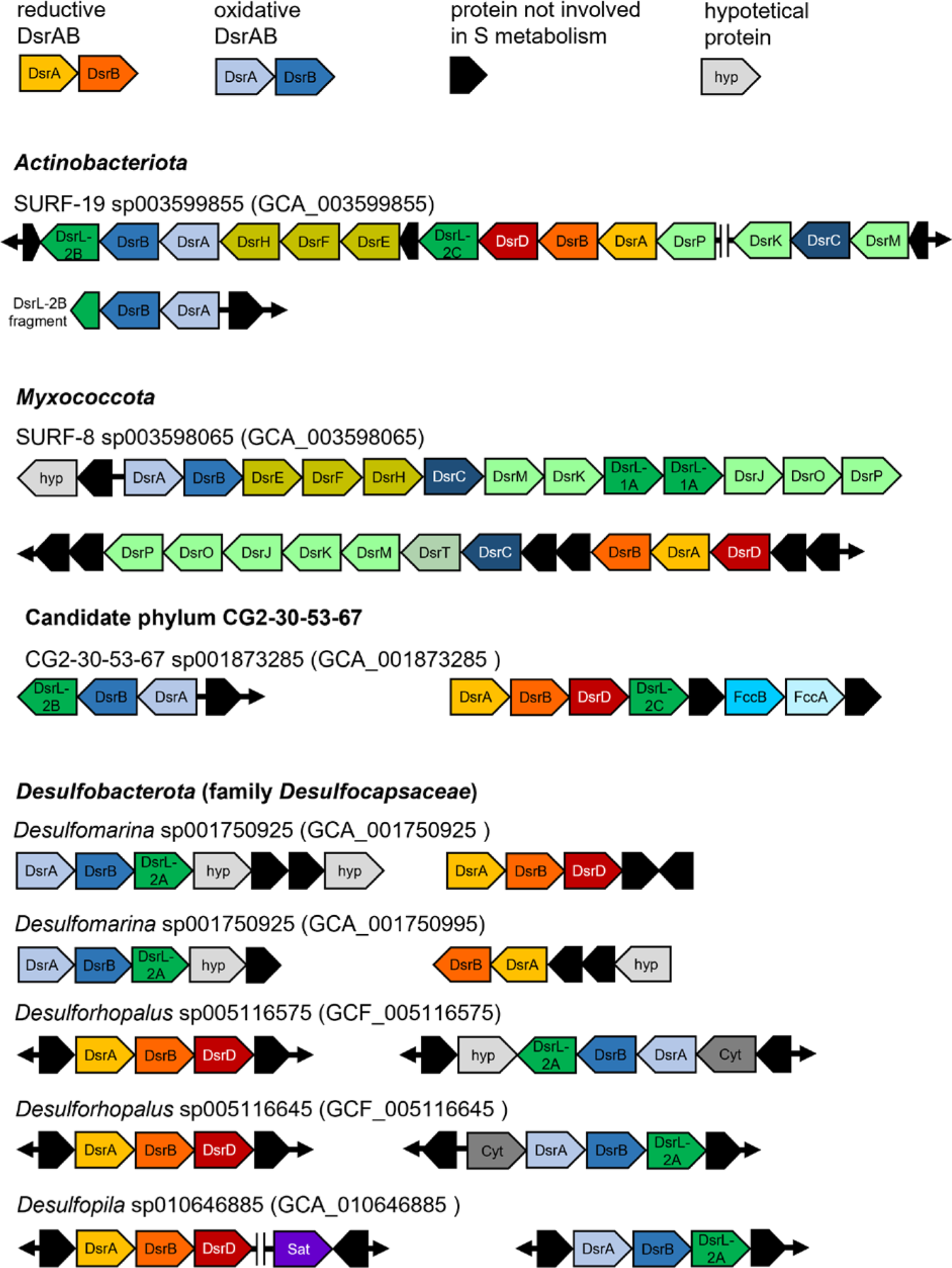
Organization of *dsr* gene clusters in MAGs encoding both a reductive and oxidative set of bacterial-type DsrAB.

The five unusual MAGs from the family *Desulfocapsaceae* within the *Desulfobacterota* were spread over the three genera *Desulforhopalus*, *Desulfomarina*, and *Desulfopila* (Fig. 4, Table S1). A uniting feature of these MAGs was their recovery from oxic-anoxic transition zones in marine surface sediments, either at methane seeps or tidal sediments. A second uniting feature was the genomic organization of the different *dsrAB* sets. Reductive *dsrAB* was always encoded upstream of *dsrD* and oxidative *dsrAB* always upstream of *dsrL*-2 (Fig. 4), which clustered together with *dsrL*-2 of canonical SOM of the genus *Chlorobium* (Fig. 3). Both *dsrAB* sets were always recovered on separate contigs and did not form operons with other *dsr*-pathway genes such as *dsrMKJOP*, *aprAB*, *qmoABC*, and *sat*. Here, a switch of the direction of dissimilatory sulfur metabolism might be regulated by differentially expressing DsrABD or DsrABL-2. Interestingly, three additional *Desulfobacterota* MAGs affiliated to the provisional genus UBA2156 encoded oxidative *dsrAB* only again along with *dsrL*-2, which clustered with *dsrL*-2 of canonical SOM of the genus *Chlorobium* (Fig. 3). Since one of these MAGs also encoded *dsrD*, it remains unclear whether reductive *dsrAB* were missed by the assembly and binning process. Vice versa, a misassembly or false binning cannot be excluded for any of the above-mentioned MAGs carrying unusual *dsrAB* combinations. However, all these MAGs were of high quality with estimated contaminations ranging from 0.7% to 6.4% (3.0 ± 2.0%, average ± standard deviation) and estimated completeness ranging from 77% to 99% (92 ± 8%, average ± standard deviation). The combination of this high binning quality with the recovery of such unusual *dsrAB* gene combinations in multiple studies from multiple environments and multiple phylogenetic lineages makes the likelihood quite high that these MAGs represent true microorganisms awaiting discovery using cultivation approaches.

Based on the findings described above, we propose to further subdivide the DsrL-2 cluster into three phylogenetically distinct subclusters to guide genome annotations. Subclusters DsrL-2A and DsrL-2B encompass (i) canonical SOM of the *Bacteroidota* and *Pseudomonadota*, (ii) *Nitrospinota* and *Nitrospirota* MAGs with an indicated oxidative sulfur metabolism, (iii) and MAGs encoding both oxidative and reductive DsrAB. For the latter group, DsrL-2A and DsrL-2B were always encoded just downstream of oxidative *dsrAB* being part of the same operon (Fig. 4). Therefore, we propose *dsrL*-2A and *dsrL*-2B to function as an indicator for an oxidative sulfur metabolism if encoded in close proximity to oxidative dsrAB. In contrast, subcluster DsrL-2C encompasses all MAGs with an indicated reductive sulfur metabolism as evidenced by the presence of DsrD and reductive DsrAB as well as absence of oxidative DsrAB. Here, we propose *dsrL*-2C to function as an indicator for a reductive sulfur metabolism if encoded in close proximity to reductive *dsrAB*. DsrL-1 subclusters A and B were already defined by Löffler et al., 2020 and encompass canonical SOM of the *Alpha*- and *Gammaproteobacteria* for subcluster DsrL-1A and canonical SOM of the *Chlorobia* and *Magnetococcia* as well as MAGs with an indicated oxidative sulfur metabolism for subcluster DsrL-1B.

**Figure 5.**
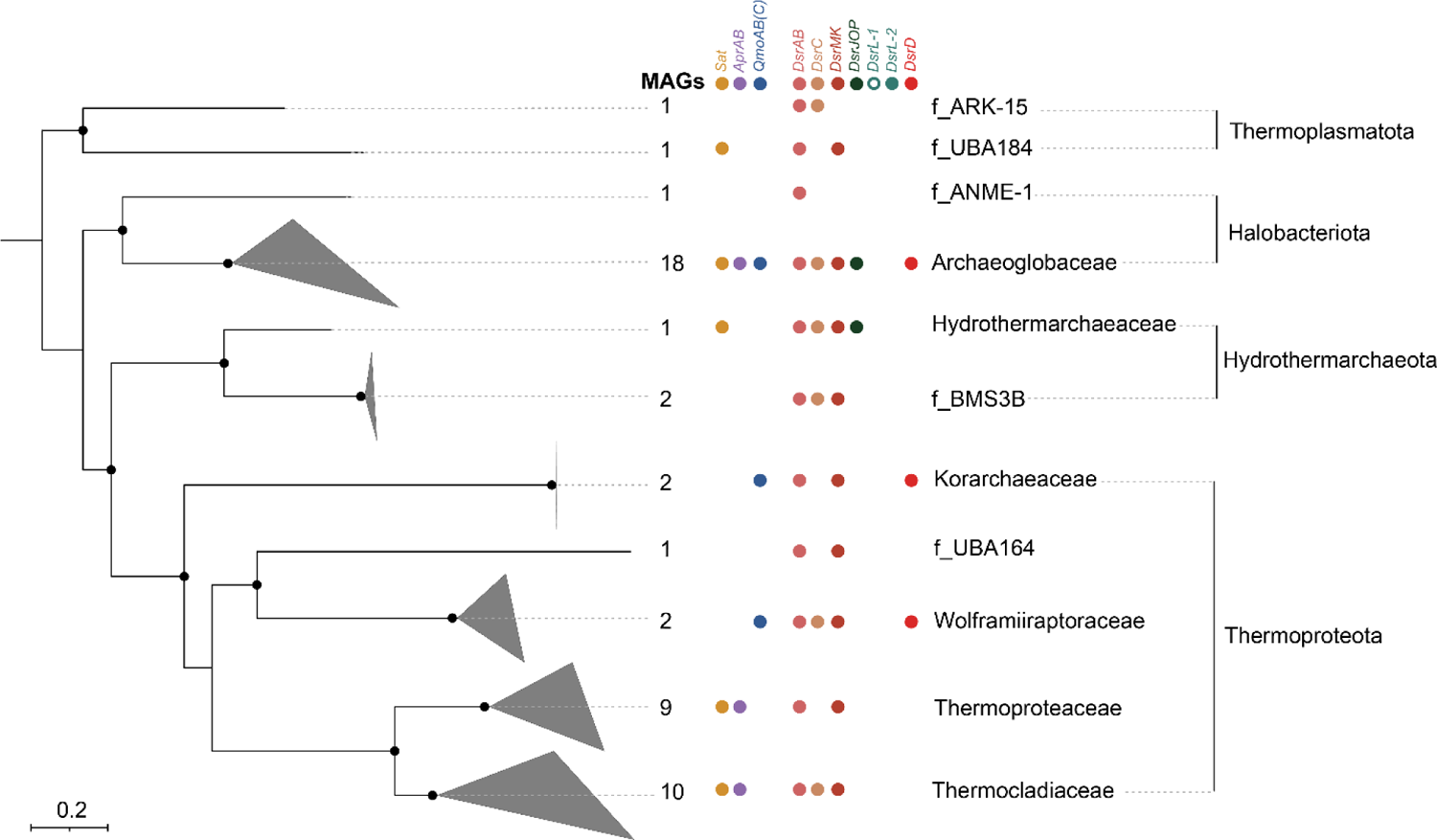
Phylogeny and *dsr*-pathway composition of *dsrAB*-encoding archaea. Archaeal genome tree inferred from 122 concatenated proteins as based on the GTDB taxonomy (Rinke et al., 2021). The phylogenetic tree was inferred from 48 archaeal (metagenome assembled) genomes. The scale bar indicates 20% sequence divergence. The maximum likelihood tree was constructed with IQ-TREE (Nguyen et al. 2015) using automatic substitution model selection (LG+F+R6) and ultrafast bootstrap analysis (n=1000). Bootstrap support is indiacted by black dots (≥90%) or black circles (70–90%). Besides *dsrAB*, the presence of *dsr*-pathway genes was indicated if >30% of *dsrAB*-containing genomes encoded the respective genes as inferred by an automated pHMM search (Zhou et al. 2022).

Archaea encoding the *dsr-*pathway are currently characterized by a solely reductively operating *dsr*-pathway. Among the 48 archaeal genomes studied, 67% belonged to cultured representatives with a known sulfate-, sulfite-, or thiosulfate-reducing metabolism within the phyla *Thermoproteota* (members of the genera *Caldivirga*, *Thermoproteus*, *Thermocladium*, *Vulcanisaeta*, and *Pyrobaculum*) and *Halobacteriota* (*Archaeoglobus* spp.). The remaining MAGs expanded the diversity of archaea encoding a *dsr-*pathway to include two additional phyla (*Hydrothermarchaeota* and *Thermoplasmatota*) and four additional families in the *Thermoproteota* (three families) and *Halobacterota* (one family). Most of the *dsrAB*-encoding archaea either represented cultured thermophiles or their MAGs were retrieved from high-temperature environments (e.g., hot springs, hydrothermal vent fluids) including their deposits (e.g., deep sea hydrothermal vent field site). The only exceptions were two *Halobacteriota* MAGs (GCA_002507545, GCA_002494625) and one *Thermoplasmatota* MAG (GCA_002503985) assembled from marine sediment or soil metagenomes, respectively (Parks et al., 2017). Notably, at least two of the families encoding the *dsr*-pathway (*Korarchaeaceae*, phylum *Thermoproteota; Archaeoglobaceae*, phylum *Halobacteriota*) were represented by MAGs (McKay et al. 2019, Wang et al. 2019) that additionally encode the complete pathway for (reverse) methanogenesis. Based on these findings, it was postulated that anaerobic methane oxidation coupled to sulfate/sulfite reduction might operate also in single microorganisms (McKay et al. 2019, Wang et al. 2019) as opposed to the standard model of syntrophic associations (Knittel and Boetius 2009).

### Genome-centric metagenomics anchors and expands *dsrAB*-based functional and taxonomic assignment

Approaches based on *dsrAB* sequence analysis have been extensively used to study the ecology of SRM and in part also SOM. These surveys were based on the assumption that there is a clear phylogenetic separation between DsrAB present in archaea with a reductive sulfur metabolism, bacteria with a reductive sulfur metabolism, and bacteria with an oxidative sulfur metabolism. We used the expanded diversity of *dsr*-pathway encoding microorganisms described above along with the indicated reductively or oxidatively operating direction of their sulfur metabolism to explore the phylogeny of the DsrAB sequence space. Our analysis showed that the distinction of archaeal reductively-operating DsrAB, bacterial reductively-operating DsrAB, and bacterial oxidatively-operating DsrAB still largely holds true. While archaeal reductively-operating DsrAB and bacterial oxidatively-operating DsrAB formed monophyletic clusters in our analysis (with the exception of laterally acquired *dsrAB*, see below), bacterial reductively-operating DsrAB were spread between these two clusters in a bush-like manner (Fig. 6). This is consistent with a recent phylogenetic analysis of DsrAB including already parts of the novel *dsrAB* sequence space discovered in MAGs from various habitats (Anantharaman et al. 2018). However, it does not reproduce anymore the monophyletic separation of reductively-operating DsrAB observed in phylogenetic analyses based mainly on canonical SRM and PCR-derived *dsrAB* sequences of environmental studies (e.g., Müller et al. 2015). Nevertheless, the distinction of reductive bacterial-type DsrAB in our (metagenome assembled) genome census was not only anchored by canonical sulfate/sulfite-reducing bacteria but contained in addition exclusively phyla whose representative MAGs encoded *DsrD* (in combination with or without *DsrL* of type 2C) strongly indicating a reductive-type sulfur metabolism as well. The only exception were the few MAGs encoding both reductive and oxidative bacterial-type DsrAB. However, also in these MAGs *dsrD* was always encoded downstream of reductive bacterial-type *dsrAB* (Fig. 4). Furthermore, the demarcation to oxidative bacterial-type DsrAB was well supported by the latest diverging reductive DsrAB cluster that contained DsrAB of *Desulfurella amilsii* (*Campylobacterota*) as an organism capable of growth by thiosulfate reduction (Fig. 6).

**Figure 6.**
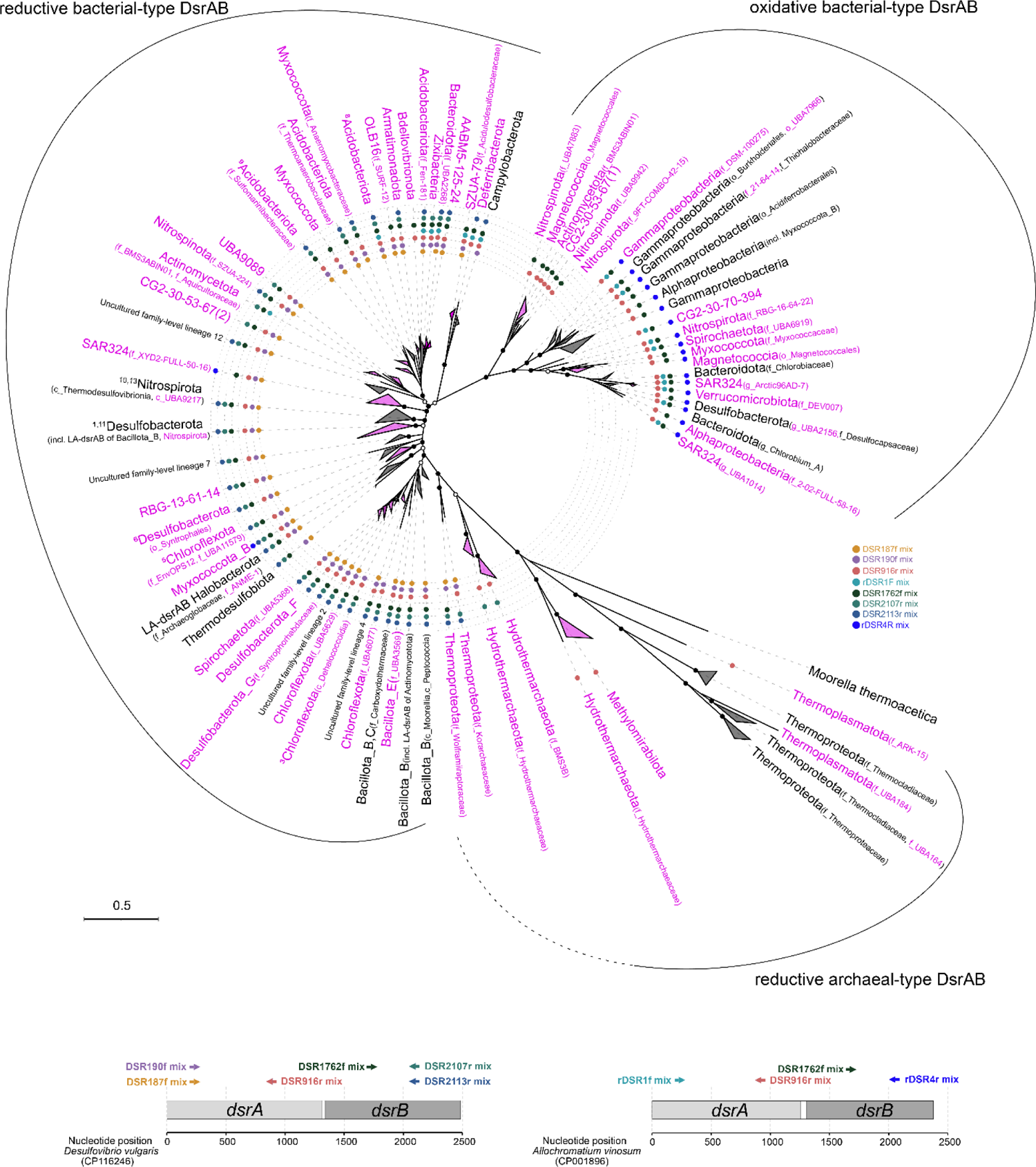
Maximum likelihood phylogeny of DsrAB sequences derived from (metagenome assembled) genomes and environmental surveys. Clades represented by a majority of *dsrAB*-encoding (metagenome assembled) genomes not affiliated to canonical SRM or SOM are shown in magenta. The coverage of inferred phylogenetic clades by published broad-range PCR primers (≥75% of sequences in a clade; 1 mismatch allowed) is indicated by colored dots. The binding positions of the evaluated primers is indicated at the bottom using *dsrAB* of *Desulfovibrio vulgaris* or *Allochromatiom vinosum* as model organism of *dsrAB* primers designed to target reductive and oxidative bacterial-type *dsrAB*, respectively. The maximum likelihood tree was constructed using deduced DsrAB amino acid sequences with IQ-TREE (Nguyen et al. 2015) using automatic substitution model selection (LG+R10) and ultrafast bootstrap analysis (n=1000). Bootstrap support is indicated by black dots (≥90%) or black circles (70–90%). The tree was inferred from 613 representative DsrAB sequences with an indel filter covering 571 amino-acid positions: 346 representative DsrAB sequences were taken from a curated DsrAB database including 7,921 pure culture and environmental sequences (as based on Müller et al. 2015) and amended with 267 DsrAB sequences derived from (metagenome assembled) genomes representing novel phylogenetic clades. Scale bar indicates 50% sequence divergence. Clades containing taxonomically resolved uncultured family-level DsrAB lineages are indicated by a superscript number based on the following denomination: 1, uncultured family-lineage 1; 3, uncultured family-lineage 3; 5, uncultured family-lineage 5; 6, uncultured family-lineage 6; 8, uncultured family-lineage 8; 9, uncultured family-lineage 9; 10, uncultured family-lineage 10; 11, uncultured family-lineage 11; 13 uncultured family-lineage 13. Please note that the numbers in brackets behind candidate phylum CG2-30-53-67 represent the two diverging *dsrAB* copies encoded by the single MAG representing this phylum. LA-*dsrAB*, laterally acquired *dsrAB*.

Previous PCR-based *dsrAB* studies described thirteen lineages of reductive bacterial-type DsrAB that represented approximate family level groups of uncultured microorganisms that could not be assigned to known taxa (Pester et al. 2012, Müller et al. 2015). Members of some of these groups were identified as abundant and active in different marine and freshwater habitats (e.g., Pester et al. 2012, Müller et al. 2015, Pelikan et al. 2016, Wörner and Pester, 2019a). Based on previous findings and the phylogenomic survey of this study, we summarize the currently known taxonomic classification of these uncultured family-level DsrAB lineages (Fig. 6, Table 1). DsrAB sequences of uncultured lineages 3 and 5 were found in members of the *Chloroflexota*. Lineage 3 members were represented by a single-cell amplified genome (SAG) recovered from deep marine subsurface sediments (Wasmund et al. 2016). Because of the low coverage of this SAG (46%, Wasmund et al. 2016), it was not part of our genome collection but was considered in our DsrAB analysis (Fig. 6, class *Dehalococcoidia*). *Chloroflexota* representing uncultured family lineage 5 were recovered from hydrothermal sediments and a bioreactor (MAG collection of Parks et al. 2017 and Zhou et al. 2020). Uncultured family lineage 6 belongs to the *Desulfobacterota* (order *Syntrophales*) and DsrAB sequences of uncultured lineages 8 and 9 were found in MAGs of terrestrial and marine *Acidobacterota*, respectively (Hausmann et al. 2018, Flieder et al. 2021). Sequences of DsrAB lineages 10 and 13 have been uncovered in *Nitrospirota* genomes from an aquifer system (Anantharaman et al. 2016). Furthermore, uncultured family lineages 1 and 11 cluster within the *Desulfobacterota.* While uncultured family lineages 1 still has an unresolved family affiliation, uncultured family lineage 11 grouped next to laterally acquired *dsrAB* of *Nitrospirota* affiliated to provisional family SM23-35. In summary, the affiliation of 8 of the 13 uncultured family-level DsrAB lineages could be resolved using (meta-)genome targeted approaches. However, the affiliation of uncultured family lineages 2, 4, 7, and 12 still awaits its discovery.

**Table 1.**
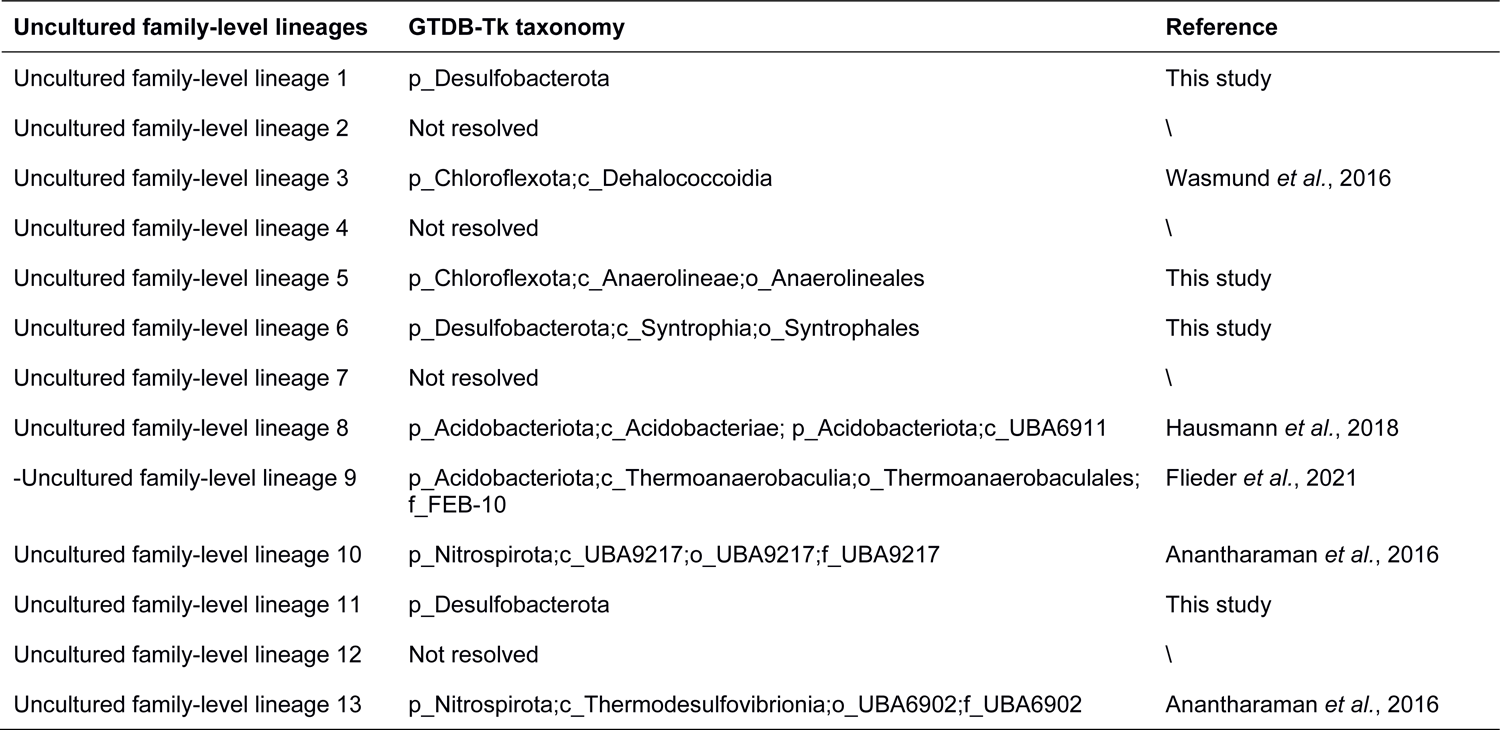
Overview of uncultured family-level DsrAB lineages as proposed by Müller et al. (2015) and their corresponding GDTB-Tk taxonomy.

Several studies have provided evidence that the distribution of *dsrAB* among extant microorganisms is represented by a combination of divergence through speciation, functional diversification and lateral gene transfer (LGT) (Klein et al. 2001, Zverlov et al. 2005, Loy et al. 2008, Müller et al. 2015, Anantharaman et al. 2018). Well documented examples are the laterally acquired *dsrAB* of a group of *Desulfotomaculum* spp. (*Bacillota*) from *Desulfobacterota* (Klein et al. 2001, Zverlov et al. 2005) or the bacterial origin of reductive *dsrAB* in members of the archaeal genus *Archaeoglobus* (*Halobacteriota*) (Müller et al. 2015). Based on our extended analysis, we conclude that 14 major taxa of *dsr*-pathway encoding microorganisms likely acquired *dsrAB* in multiple lateral gene transfer events. These encompass besides the *Bacillota* and *Halobacterota* also the *Methylomirabilota, Chloroflexota*, *Alphaproteobacteria*, *Magnetococcia*, *Hydrothermarchaeota*, *Desulfobacterota*, *Actinomycetota*, *Nitrospirota*, *Nitrospinota*, *Bacteroidota*, *Spirochaetota*, *Myxococcota*, and candidate phyla CG2-30-53-67 and SAR324 (Fig. S1). The latter seven are especially interesting because they harbor members with either reductive *dsrAB*, oxidative *dsrAB*, or both (Fig. 6, Fig. S1). Analysis of our extended *dsrAB* dataset could not reproduce the postulated LGT of bacterial *dsrAB* encoded to archaea of the families *Wolframiiraptoraceae* (previously referred to as Aigarchaota pSL4) and *Korarchaeaceae* (*Ca.* Methanodesulfokores washburnensis) (Müller et al. 2015, McKay et al. 2019) despite them showing higher similarity to reductive bacterial-type DsrAB of *Bacillota* than to reductive DsrAB of all other archaea (Fig. 6). More in-depth phylogenetic studies will have to show whether lateral gene transfer of *dsrAB* occurred in *Wolframiiraptoraceae* and the *Korarchaeaceae* as well.

With the updated *dsrAB* database provided in this review, *dsrAB*-based marker gene surveys will greatly benefit as sequences can be better taxonomically anchored. To this end, we provide an updated *dsrA* and *dsrB* gene reference database including the sequences from 902 bacterial and 48 archaeal genomes and MAGs analyzed in this study (available under www.arb-silva.de/projects/), which will be useful for *dsrAB* amplicon sequencing analyses (e.g., Müller et al. 2015, Pelikan et al. 2016, Vigneron et al. 2018, Wörner and Pester, 2019b). We further evaluated the coverage of those genomes and MAGs with primer sets designed to target reductive bacterial-type (Pelikan et al. 2016) and oxidative bacterial-type (Loy et al. 2009, Müller et al. 2015) *dsrA* and *dsrB* sequences (Table 2 and Tables S2-S4). A good coverage of near full-length reductive bacterial-type *dsrAB* can be achieved using the primers DSR190f mix (88%) and DSR2107r mix (88%) for the great majority of (putative) SRM (Fig. 6 and Table 2, for details see Tables S3-S4). For amplicon sequencing, we can confirm the recommendation of Pelikan et al. (2016) to use the primer pair DSR1762f mix and DSR2107r mix, which cover most (97% and 88%, respectively) of the reductive bacterial-type *dsrB* (Fig. 6, Table 2). However, for extended coverage of the few MAGs within the *Bdellovibrionota*, *Campylobacterota*, *Deferribacterota*, SAR324, SZUA-79, OLB16, or AABM5-125-24 new primer variants will need to be designed for both near-full length and short *dsr(A)B* amplicons (Table 2). Primer pairs rDSR1f mix and rDSR4r mix designed to amplify near full-length oxidative bacterial-type *dsrAB* (Loy et al. 2009) have an acceptable coverage (68% and 95%, respectively), but do not cover a considerable fraction of the *dsrAB*-encoding *Alphaproteobacteria, Gammaproteobacteria*, and *Magnetococcia* as well as oxidative *dsrAB* within the *Nitrospinota*, *Nitrospirota*, and candidate phylum CG2-30-53-67. However, the majority of oxidative bacterial-type *dsrB* sequences is covered by the primer pairs DSR1762f mix (94%) and rDSR4r mix (95%) for direct short amplicon sequencing (Table 2 and Tables S3-S4). No primers were published so far to specifically amplify reductive archaeal-type *dsrAB*.

**Table 2.**
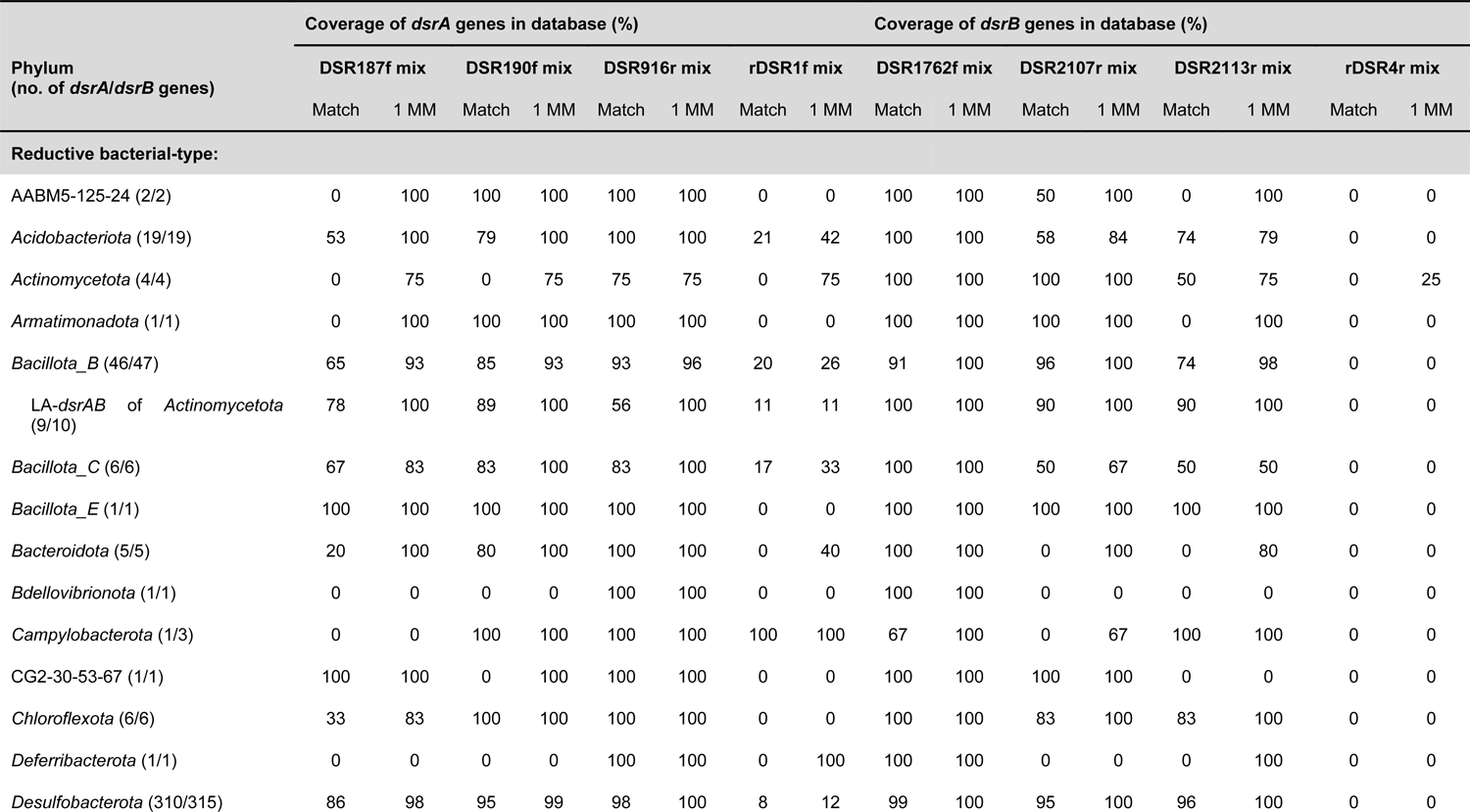
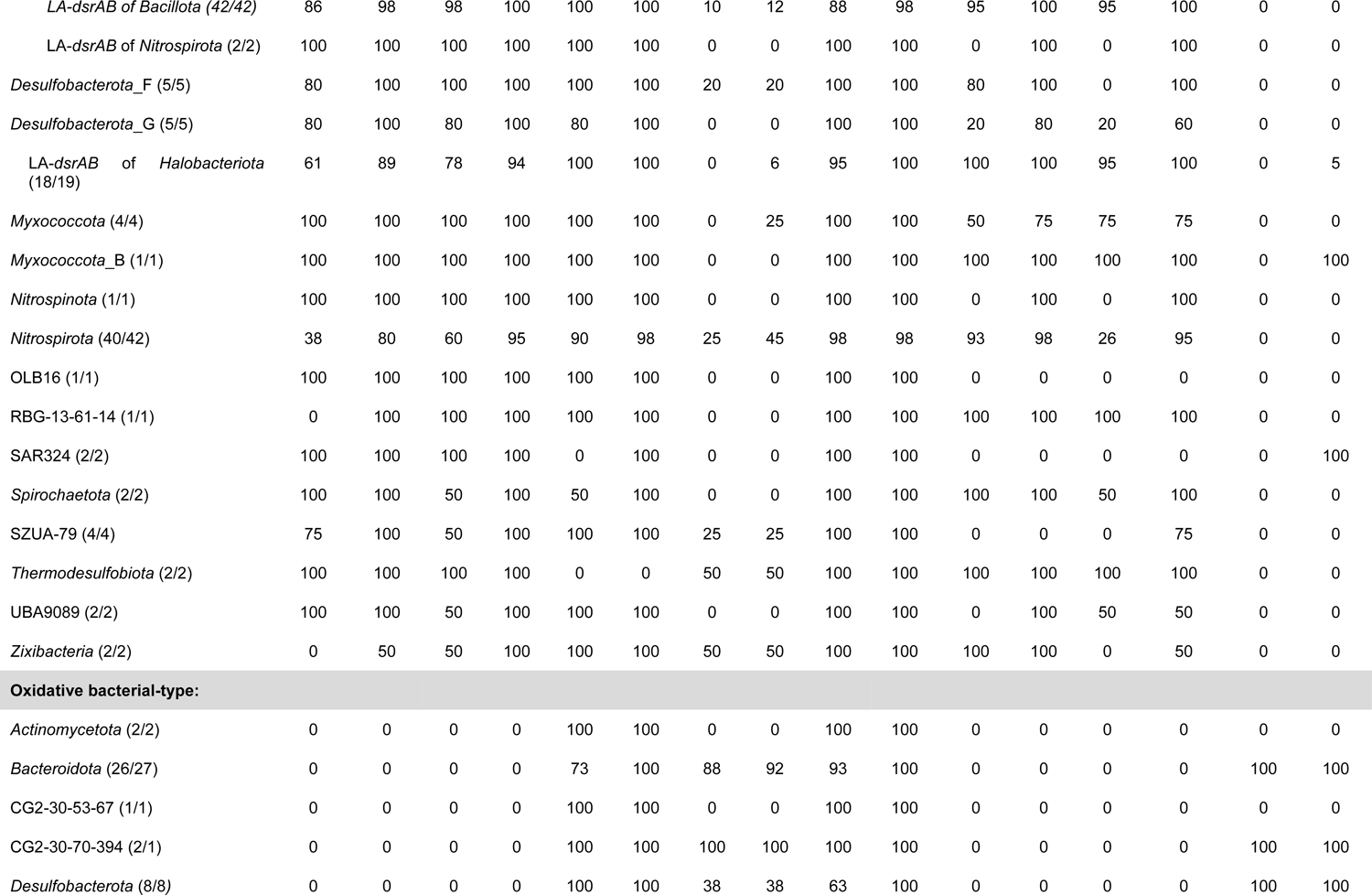
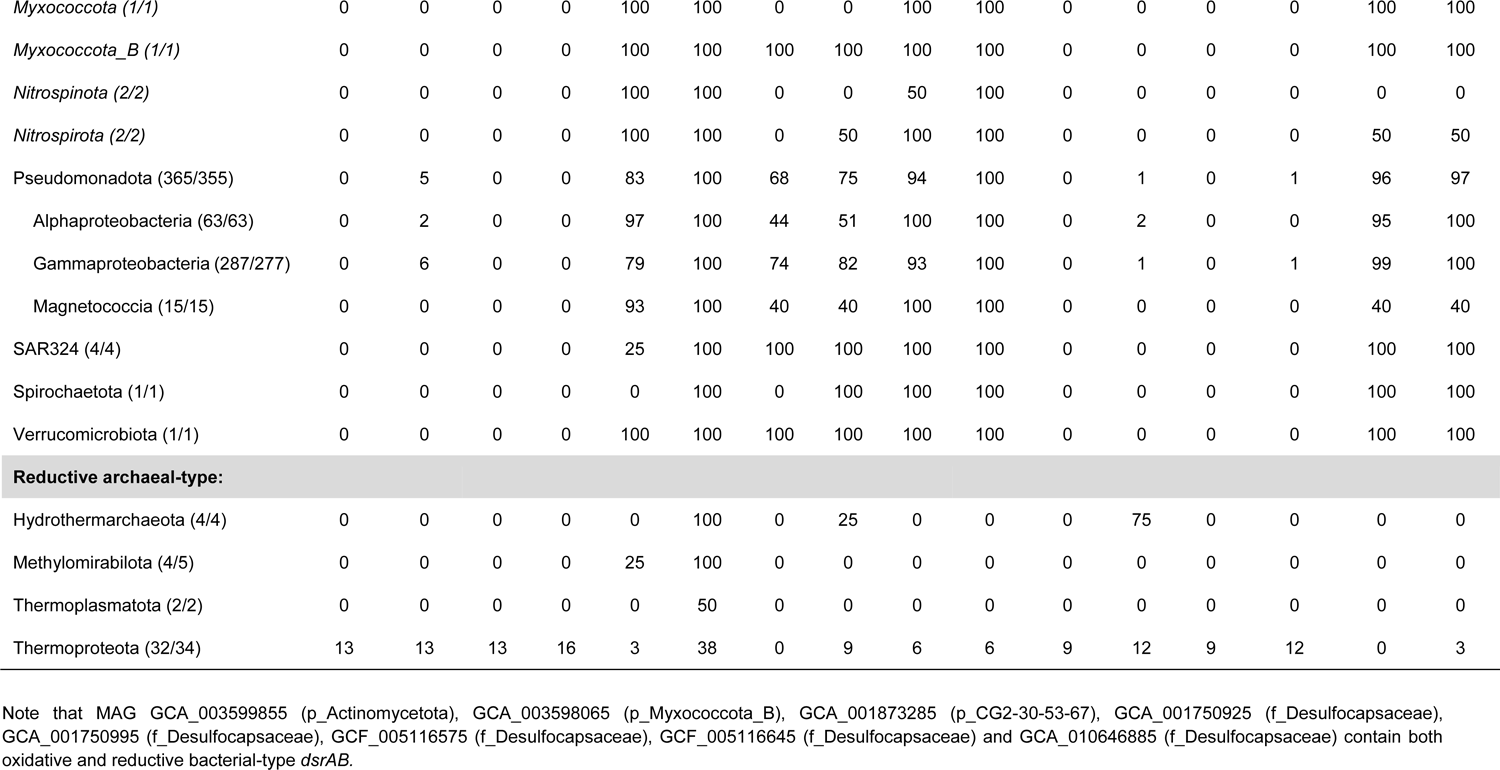
Coverage of *dsrAB* primers targeting reductive (DSR mixes) or oxidative (rDSR mixes) bacterial-type *dsrAB*. Coverages are provided as full match or with the tolerance of one mismatch (1 MM). The number of analyzed *dsrA* and *dsrB* sequences are indicated behind the respective phyla and were depending on the completeness of the respective gene sequences at the primer binding sites. If phyla encompassed members with reductive and/or oxidative *dsrAB* they were listed along with the representing lower taxonomic ranks.

### Insights into the ecophysiology of newly discovered SRM

For most of the newly discovered microorganisms encoding a *dsr*-pathway only a (partial) genome sequence is available so far. Even though their genomic context provides clues about a reductively or oxidatively operating dissimilatory sulfur metabolism, we still miss a large part of their actual physiology. Efforts to enrich and isolate these microorganisms into culture will remain the best approach to understand their biology but will also take time. An alternative to cultivation is to understand the ecophysiology of these newly discovered SRM (and SOM) in their natural setting using controlled experimental setups along with studying their activity responses at the transcriptome and/or proteome level or using isotope labeling techniques at the population or single-cell level. In the following, we describe several examples, where this has been achieved.

*Acidobacteriota* encoding a *dsr*-pathway were first discovered in pristine low-sulfate environments including peatlands (Hausmann et al. 2018) and aquifers (Anantharaman et al. 2018) but also in acidic sulfide mine waste rock sites (Anantharaman et al. 2018) and later in marine surface sediments (Flieder et al. 2021). Clone library-based studies targeting *dsrAB* extend the habitat range to deep marine sediments below the sulfate-methane transition zone and high-temperature environments (reviewed in Pester et al. 2012, Müller et al. 2015). A few studies succeeded to investigate the encoded metabolic potential, the transcriptional activity, as well as the abundance and distribution of *dsr*-pathway encoding *Acidobacteriota* in more detail.

Peatland *Acidobacteriota* encoding a *dsr*-pathway were studied in detail in a small acidic fen in the Fichtel mountains located in Central Europe. Here, they make up roughly two thirds of microorganisms encoding reductively operating bacterial *dsrAB* (Pelikan et al. 2018) and contribute a considerable fraction to the overall microbial peat soil transcriptome (>2% of all mRNA reads; Hausmann et al. 2018), implying a predominant role in the hidden sulfur cycle of this habitat. They are affiliated to four different families (*Acidobacteriaceaea*, SBA1, *Bryobacteraceae*, and UBA7540) within the class *Terriglobia* (comprising former uncultured *dsrAB* family-level lineage 8) with some recovered MAGs encoding the full canonical pathway of sulfate reduction while others harboring only genes for sulfite reduction. The latter encoded in addition enzymes that can liberate sulfite from organosulfonates, implying organic sulfur compounds as complementary energy sources. Interestingly, these *Acidobacteriota* encoded also the full respiratory chain for aerobic respiration including low and high affinity terminal oxidases as well as a large enzymatic repertoire for polysaccharide degradation and sugar utilization. In addition, capabilities for a fermentative lifestyle and hydrogen oxidation were encoded as well. This “Swiss army knife”-array of potential energy metabolism variants opens up a lot of possibilities how these *dsr*-pathway encoding *Acidobacteriota* may cope with the fluctuating redox conditions in peat soils. The redox state of these typically water-saturated soils can change dramatically and is mainly driven by changes in the water table through rainfall and droughts. In addition, lateral flow of water can heavily influence the topography of redox gradients in peat soils through space and time (Frei et al. 2012, Pester et al. 2012). In response, *dsr*-pathway encoding *Acidobacteriota* where postulated to be able of switching from a sulfate-reducing or, in case of sulfate shortage, fermentative lifestyle under anoxic conditions to aerobic respiration under oxic conditions using polysaccharides or low-molecular weight organic compounds as substrates. Especially the potential use of polysaccharides under sulfate reducing conditions would differentiate them from canonical SRM, which are not able to degrade organic polymers (Rabus et al. 2013). Also a reversal of the *dsr*-pathway for sulfur oxidation in combination with aerobic respiration was proposed (Hausmann et al. 2018), although the genomic context suggests rather a reductively operating sulfur metabolism (see above).

Peat soil incubations under controlled substrate and sulfate supply may provide insights into the postulated metabolism of *dsrAB*-encoding *Acidobacteriota*. When peat soil was incubated anoxically with individual fermentation intermediates (formate, acetate, propionate, lactate or butyrate) with and without externally supplied sulfate, *dsr*-pathway encoding *Acidobacteriota* showed a steady transcriptional activity including all genes of the *dsr-*pathway. However, there was no significant increase of transcriptional activity triggered by either one of the individually supplied low-molecular weight compounds indicating that the activity of the respective *Acidobacteriota* rather relied on organic substances already present in the peat itself (Hausmann et al. 2018). Extending upon these initial results, Dyksma and Pester (submitted) incubated peat soil in a bioreactor setting under alternating oxic (50% air saturation) and anoxic conditions and a steady supply of pectin as an abundant terrestrial plant polysaccharide. Indeed, a *dsr*-pathway encoding Acidobacterium differentially expressed the full canonical pathway of sulfate reduction under anoxic conditions and the full respiratory chain under oxic conditions providing experimental evidence that facultatively anaerobic SRM within the *Acidobacteriota* exist (Dyksma and Pester, submitted). Similar results were already indicated in studies on model SRM within the *Desulfobacterota*, i.e. *Desulfovibrio* species, albeit at much lower oxygen concentrations. *Desulfovibrio* spp. typically encode high-affinity bd-type terminal oxidases only, which are implied to function in oxygen detoxification rather than aerobic growth (Ramel et al. 2015). When grown in semi-solid media within an oxygen gradient, *Desulfovibrio magneticus* formed in the absence of sulfate a visible band at the oxic-anoxic interface and the authors interpreted this as micro-oxic growth coupled to oxygen respiration (Lefèvre et al. 2016). In a more detailed study, a strain of *Desulfovibrio vulgaris* Hildenborough was exposed to O_2_-driven laboratory adaptive evolution and acquired via point mutations as well as deletions/insertions the ability to gain energy from oxygen respiration under microoxic conditions (0.65% O_2_, Schoeffler et al. 2019). Since the enzymatic systems required for both sulfate and oxygen respiration were already present in the genome of *D. vulgaris*, only a limited number of mutations were apparently required to redirect the flow of reducing equivalents towards aerobic respiration coupled to growth (Schoeffler et al. 2019).

Marine *Acidobacteriota* were first indicated in *dsrAB*-based surveys (Müller et al. 2015) and recently their respective genomes could be recovered from marine surface sediments in the Arctic off the coast of Svalbard (Flieder et al. 2021). Here, they comprised the second most abundant *dsrAB*-encoding phylum after the *Desulfobacterota* (on average 13%) and represented 4% of *dsrB* transcripts emphasizing their *in situ* activity. When expanded to a global marine *dsrAB* dataset, acidobacterial *dsrAB* averaged 15% in marine sediments worldwide (Flieder et al. 2021). They are affiliated to a different class (*Thermoanaerobaculia;* family FEB-10) than peatland *dsr*-pathway encoding *Acidobacteriota* and comprise former uncultured *dsrAB* family-level lineage 9. Detailed annotation of their genomes revealed the metabolic potential for various respiratory pathways based on oxygen, nitrous oxide, metal-oxide, tetrathionate, sulfur and sulfate/sulfite as terminal electron acceptor. The potential for sulfur disproportionation was indicated as well. Potential electron donors comprised cellulose, proteins, cyanophycin, hydrogen, and acetate (Flieder et al. 2021). In summary, both terrestrial and marine *dsr*-pathway encoding *Acidobacteriota* likely represent an ecologically important but so far overlooked group of SRM with a large metabolic versatility in respect to potential substrates including organic polymers and alternative electron acceptors including oxygen. This strategy might be key for their success to cope with fluctuating conditions, e.g., during drying and rewetting in peatland soils, varying oxygen penetration during tidal cycles in coastal sediments and at oxic-anoxic transition zones in freshwater and marine sediments. This metabolic flexibility is also a major hallmark differentiating them from canonical SRM, mainly within the *Desulfobacterota* and *Bacillota*, giving them a competitive advantage in environments with highly variable redox conditions.

The metabolic flexibility to switch between sulfate reduction and aerobic respiration was also indicated in metagenomic and metatranscriptomic surveys of microbial mat-inhabiting members of the *Bacteroidota* family UBA2268 (Kapabacteria). In contrast to their well-studied phototrophic and sulfur-oxidizing relatives within the *Chlorobiaceae*, UBA2268-related MAGs retrieved from microbial mats of hot springs or groundwater encode reductive *dsrAB* as well as *dsrD* and *dsrL* of type 2C. For one of these MAGs (*Candidatus* Thermonerobacter thiotrophicus), the genome was annotated in greater detail and its transcriptional profile characterized during the diel cycle in the microbial mat of the thermal outflow of Mushroom Spring in Yellowstone National Park, USA (Thiel et al. 2019). Despite being a low-sulfate environment (<200 µM), the phototrophic microbial mat was characterized by high sulfate reduction rates (>5 µmol cm^-3^ d^-1^) during the night, which ceases during daytime because of oxygen production by cyanobacteria-driven photosynthesis (Dillon et al. 2007). Accordingly, *Ca*. T. thiotrophicus showed strong expression of all *dsr*-pathway genes during the night and a sharp decrease in its transcript levels during daytime. *Ca*. T. thiotrophicus also encoded a full respiratory chain including alternative complex III, an aa_3_-type low-affinity terminal oxidase as well as a bd-type high-affinity terminal oxidase. Interestingly, genes encoding the aa_3_-type low-affinity terminal oxidase were differentially expressed as compared to *dsr*-pathway genes. Their highest expression levels were observed at light-dark transitions in the morning and evening (Thiel et al. 2019) corresponding to increasing and decreasing oxygen levels in the mat, respectively, but avoiding times of oxygen (over)saturation during daytime (Dillon et al. 2007), when genes encoding oxidative stress response dominated the transcriptional profile (Thiel et al. 2019). In contrast, genes encoding the bd-type terminal oxidase showed highest expression during the night implying a role in oxygen detoxification at low oxygen levels during active sulfate reduction. The absence of encoded CO_2_-fixation pathways and the increased expression of genes involved in glycolysis/gluconeogenesis, the TCA cycle, and acetate-related metabolism during the night indicated a heterotrophic lifestyle based on small organic molecules, which primarily correlated with sulfate reduction (Thiel et al. 2019).

Another intriguing group are the many *dsrAB*-encoding *Nitrospirota* members with an indicated reductive sulfur metabolism, which were encountered in environments of mainly moderate temperatures and that are distinct from their thermophilic, sulfate-reducing relatives within the genus *Thermodesulfovibrio*. When excluding *Thermodesulfovibrio* spp., 37 additional MAGs encoding an indicated reductively operating *dsr*-pathway were recovered representing 10 candidate families within the phylum *Nitrospirota*. Typically, these MAGs were recovered from low-sulfate environments encompassing rice paddy soil (Zecchin et al. 2018), permafrost soils (Woodcroft et al. 2018), freshwater sediments, aquifer sediments, groundwater, the terrestrial and marine deep subsurface (Jungbluth et al. 2017, Anantharaman et al. 2018, Probst et al. 2018). However, a few were also recovered from brackish (Arshad et al. 2017) and saline marine environments (Kato et al. 2018). Among the mesophilic, low-sulfate adapted *Nitrospirota,* representatives from rice paddies were studied in more detail. From paddy soil that was used to grow rice in the presence and absence of gypsum (CaSO_4_ × 2 H_2_O), the partial genome of *dsr*-pathway encoding *Candidatus* Sulfobium mesophilum (*Nitrospirota* family UBA6898) could be recovered. Parallel metaproteomics revealed active expression of its *dsr*-pathway under gypsum amendment in support of a sulfate-reducing lifestyle. Interestingly, *Ca*. S. mesophilum also encoded the full pathway of dissimilatory nitrate reduction to ammonia, which was expressed in the treatment without gypsum amendment. The relative abundance of *Ca*. S. mesophilum was similar under both treatments, indicating that it maintains a stable population in rice paddy soils while shifting its primary energy metabolism. In contrast to the *Acidobacteriota* described above, *Ca*. S. mesophilum was rather adapted to the breakdown of classical substrates of SRM covering the metabolic potential to utilize butyrate, formate, H_2_, and acetate as an electron donor (Zecchin et al. 2018).

The *Actinomycetota* represented yet another unusual phylum harboring *dsr*-pathway encoding members (Müller et al. 2015). Besides the unusual *Actinomycetota* MAG GCA_003599855, which encoded reductive and oxidative *dsrAB*, all other retrieved *Actinomycetota* could be split into two major groups as based on the completeness of their *dsr*-pathway and habitat preference. All members of the class *Coriobacteriia* (five genomes/MAGs within the genera *Gordonibacter*, *Rubneribacter*, *Berryella*, and UBA8131) encoded only the genetic potential to reduce sulfite to sulfide, included *dsrD*, lacked *dsrL*, and were so far isolated or encountered in intestines of humans and animals including pig, chicken, and termites (Würdemann et al. 2009, Selma et al. 2014, Medvecky et al. 2018, Parks et al. 2018, Wylensek et al. 2020). Cultured representatives from the genera *Gordonibacter* and *Berryella* are strict anaerobes supporting the notion that the encoded reductive bacterial-type *dsrAB* and the presence of *dsrD* point towards a reductive sulfur metabolism. However, dissimilatory sulfite reduction by these microorganisms still awaits experimental validation. Sulfite in the gut environment is likely derived from sulfonates, i.e. organic sulfur compounds with a SO ^2-^ moiety, such as the amino acid taurine (Wei et al. 2021) or the sugar sulfoquinovose (Hanson et al. 2021). The second major group within the *dsr*-pathway encoding *Actinomycetota* encodes the full *dsr*-pathway including *dsrD* and *dsrL* of type 2C. They represent uncultured members of the classes *Thermoleophilia* (Kato et al. 2018) and *Aquicultoria* (Jiao et al. 2021), with the latter encoding the oxygen-sensitive Wood-Ljungdahl pathway pointing towards a strictly anaerobic lifestyle and a reductively operating sulfur metabolism. In contrast to the incomplete *dsr*-pathway encoding members of the *Coriobacteriia*, they were found in terrestrial and marine environments including groundwater, the terrestrial subsurface (Jiao et al. 2021), and deep-sea massive sulfide deposits (Kato et al. 2018).

## Conclusion

Metagenome-driven discoveries have opened a new window into the hidden diversity of SRM. We can now start to appreciate that besides the four bacterial and two archaeal phyla harboring cultured SRM, the potential to perform dissimilatory sulfate/sulfite reduction extends to a total of 23 bacterial and 4 archaeal phyla. Many of the phyla now recognized to play a role in sulfur cycling were represented by *dsrAB*-encoding MAGs recovered from low-sulfate environments, supporting the notion that hidden or cryptic sulfur cycling in low sulfate environments is an understudied area. For a few of these potential SRM, such as members of the *Acidobacteriota*, mesophilic *Nitrospirota,* and *Bacteriodata* family UBA2268 (Kapabacteria), meta-omics based studies under constrained environmental conditions could provide strong evidence of a sulfate-reducing lifestyle. However, the large majority of novel, putative SRM still await experimental confirmation of their physiology. Furthermore, we could show that the primers used in *dsrAB*-based approaches cover a large fraction of the novel diversity of SRM, with many of the previously taxonomically unresolved *dsrAB* lineages now anchored by *dsrAB*-encoding MAGs. As such, *dsrAB*-based surveys can be used with confidence in the future to explore the enigmatic world of a functional microbial guild that has shaped biogeochemical cycling on Earth since the Archaean (Shen et al. 2001, Wacey et al. 2011).

## Acknowledgement

This work was supported by the Alexander von Humboldt Foundation (MD), the Leibniz Institute DSMZ (SD, EK, DKN), the German Research Foundation DFG (PE2147/3-1 to MP), the Austrian Science Fund FWF (P 31996-B to AL), and the US National Science Foundation (OCE 2049478 to KA).

## Conflict of interest

None declared.

---

**Supplementary Figure S1.**
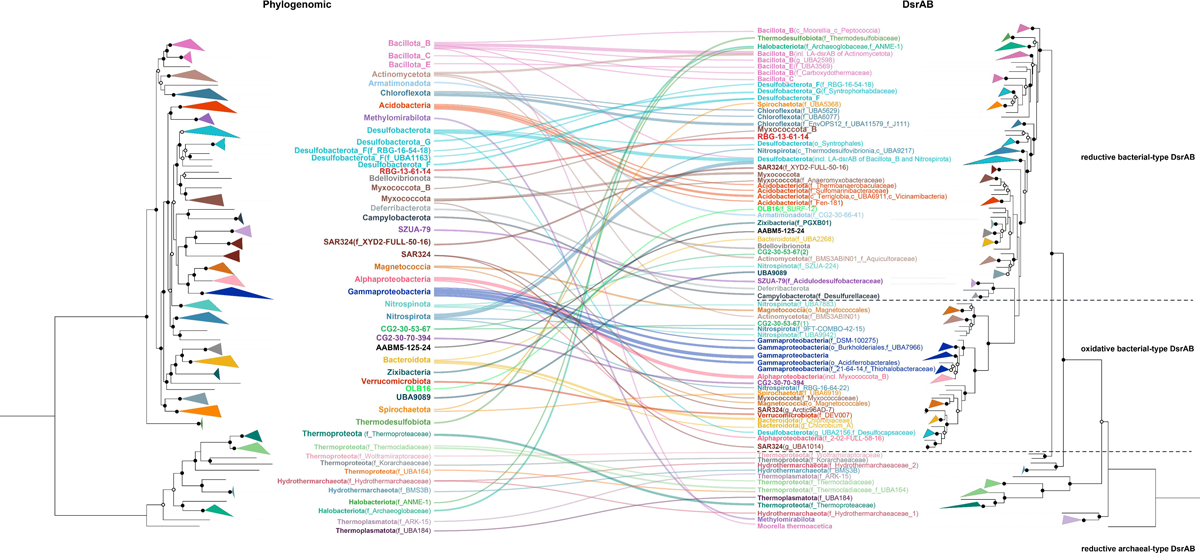
Comparison of phylogenomic and DsrAB trees for microorganisms representing all inferred *dsrAB*-encoding archaeal and bacterial lineages. The phylogenomic tree was inferred from a set of 43 conserved single-copy marker genes obtained with CheckM (Parks *et al*., 2015) using 38 representative archaeal and 207 representative bacterial (metagenome-assembled) genomes. The phylogenomic maximum likelihood tree was constructed with IQ-TREE (Nguyen *et al*. 2015) using automatic substitution model selection (LG+R10) and ultrafast bootstrap analysis (n=1000). The DsrAB tree was constructed using 269 deduced DsrAB amino acid sequences, which were extracted from 245 representative (metagenome-assembled) genomes. The DsrAB maximum likelihood tree was constructed with IQ-TREE (Nguyen *et al*. 2015) using automatic substitution model selection (LG+R8) and ultrafast bootstrap analysis (n=1000). Bootstrap support is indicated by black dots (≥90%) or black circles (70–90%).

## Notes

### Competing Interest Statement

The authors have declared no competing interest.

## References

Anantharaman K, Brown CT, Hug LA et al. Thousands of microbial genomes shed light on interconnected biogeochemical processes in an aquifer system. Nat Commun 2016;7:13219.

Anantharaman K, Hausmann B, Jungbluth SP et al. Expanded diversity of microbial groups that shape the dissimilatory sulfur cycle. ISME J 2018;12:1715–28.

Arshad A, Dalcin Martins P, Frank J et al. Mimicking microbial interactions under nitrate-reducing conditions in an anoxic bioreactor: enrichment of novel *Nitrospirae* bacteria distantly related to *Thermodesulfovibrio*. Environ Microbiol 2017;19:4965–77.

Barton LL, Ritz NL, Fauque GD et al. Sulfur cycling and the intestinal microbiome. Dig Dis Sci 2017;62:2241–57.

Boetius A, Ravenschlag K, Schubert CJ et al. A marine microbial consortium apparently mediating anaerobic oxidation of methane. Nature 2000;407:623–6.

Bowles MW, Mogollón JM, Kasten S et al. Global rates of marine sulfate reduction and implications for sub–sea-floor metabolic activities. Science 2014;344:889–91.

Breitburg D, Levin LA, Oschlies A et al. Declining oxygen in the global ocean and coastal waters. Science 2018;359:eaam7240.

Bush T, Diao M, Allen RJ et al. Oxic-anoxic regime shifts mediated by feedbacks between biogeochemical processes and microbial community dynamics. Nat Commun 2017;8:789.

Canfield DE, Farquhar J. The Global Sulfur Cycle. In: Knoll AH, Canfidld DE, Konhauser KO (eds.). Fundamentals of Geobiology. John Wiley & Sons, 2012, 49–64.

Colman DR, Lindsay MR, Amenabar MJ et al. Phylogenomic analysis of novel Diaforarchaea is consistent with sulfite but not sulfate reduction in volcanic environments on early Earth. ISME J 2020;14:1316–31.

Coskun ÖK, Özen V, Wankel SD et al. Quantifying population-specific growth in benthic bacterial communities under low oxygen using H_2_^18^O. ISME J 2019;13:1546–59.

Crowe SA, Paris G, Katsev S et al. Sulfate was a trace constituent of Archean seawater. Science 2014;346:735–9.

Dahl C. Sulfur Metabolism in Phototrophic Bacteria. In: Hallenbeck PC (ed.). Modern Topics in the Phototrophic Prokaryotes: Metabolism, Bioenergetics, and Omics. Springer, 2017, 27–66.

Dedysh SN, Sinninghe Damsté JS. Acidobacteria. eLS. Chichester: John Wiley & Sons, 2018, a0027685.

Diao M, Huisman J, Muyzer G. Spatio-temporal dynamics of sulfur bacteria during oxi-anoxic regime shifts in a seasonally stratified lake. FEMS Microbiol Ecol 2018;94:fiy040.

Diaz RJ, Rosenberg R. Spreading dead zones and consequences for marine ecosystems. Science 2008;321:926–9.

Dillon JG, Fishbain S, Miller SR et al. High rates of sulfate reduction in a low-sulfate hot spring microbial mat are driven by a low level of diversity of sulfate-respiring microorganisms. Appl Environ Microbiol 2007;73:5218–26.

Dopson M, Johnson DB. Biodiversity, metabolism and applications of acidophilic sulfur-metabolizing microorganisms. Environ Microbiol 2012;14:2620–31.

Dyksma S, Pjevac P, Ovanesov K et al. Evidence for H2 consumption by uncultured Desulfobacterales in coastal sediments. Environ Microbiol 2018;20:450–61.

Fakhraee M, Hancisse O, Canfield DE et al. Proterozoic seawater sulfate scarcity and the evolution of ocean–atmosphere chemistry. Nat Geosci 2019;12:375–80.

Fakhraee M, Katsev S. Organic sulfur was integral to the Archean sulfur cycle. Nat Commun 2019;10:4556.

Ferreira D, Barbosa ACC, Oliveira GP et al. The DsrD functional marker protein is an allosteric activator of the DsrAB dissimilatory sulfite reductase. Proc Natl Acad Sci USA 2022;119: e2118880119.

Flieder M, Buongiorno J, Herbold CW et al. Novel taxa of Acidobacteriota implicated in seafloor sulfur cycling. ISME J 2021;15:3159–80.

Florentino AP, Stams AJM, Sánchez-Andrea I. Genome Sequence of *Desulfurella amilsii* strain TR1 and comparative genomics of *Desulfurellaceae* family. Front Microbiol 2017;8:222.

Florentino AP, Pereira IAC, Boeren S et al. Insight into the sulfur metabolism of *Desulfurella amilsii* by differential proteomics. Environ Microbiol 2019;21:209–25.

Frei S, Knorr KH, Peiffer S et al. Surface micro-topography causes hot spots of biogeochemical activity in wetland systems: A virtual modeling experiment. J Geophys Res Biogeosci 2012;117:G00N12.

Frolov EN, Lebedinsky AV, Elcheninov AG et al. Taxonomic proposal for a deep branching bacterial phylogenetic lineage: transfer of the family *Thermodesulfobiaceae* to *Thermodesulfobiales* ord. nov., Thermodesulfobiia classis nov. and Thermodesulfobiota phyl. nov. Syst Appl Microbiol 2023;46:126388.

Habicht KS, Gade M, Thamdrup B et al. Calibration of sulfate levels in the Archean Ocean. Science 2002;298:2372–4.

Halevy I. Production, preservation, and biological processing of mass-independent sulfur isotope fractionation in the Archean surface environment. Proc Natl Acad Sci USA 2013; 110:17644–9.

Hanson BT, Dimitri Kits K, Löffler J et al. Sulfoquinovose is a select nutrient of prominent bacteria and a source of hydrogen sulfide in the human gut. ISME J 2021;15:2779–91.

Hausmann B, Pelikan C, Herbold CW et al. Peatland Acidobacteria with a dissimilatory sulfur metabolism. ISME J 2018;12:1729–42.

Hausmann B, Pelikan C, Rattei T et al. Long-term transcriptional activity at zero growth of a cosmopolitan rare biosphere member. mBio 2019;10:e02189–18.

Hug LA, Thomas BC, Sharon I et al. Critical biogeochemical functions in the subsurface are associated with bacteria from new phyla and little studied lineages. Environ Microbiol 2016;18:159–73.

Jenny J-P, Francus P, Normandeau A et al. Global spread of hypoxia in freshwater ecosystems during the last three centuries is caused by rising local human pressure. Glob Chang Biol 2016;22:1481–9.

Jiao JY, Fu L, Hua ZS et al. Insight into the function and evolution of the Wood-Ljungdahl pathway in *Actinobacteria*. ISME J 2021;15:3005–18.

Jørgensen BB. Sulfur biogeochemical cycle of marine sediments. Geochem Perspect 2021;10: 145–6.

Jungbluth SP, Glavina Del Rio T, Tringe SG et al. Genomic comparisons of a bacterial lineage that inhabits both marine and terrestrial deep subsurface systems. Peer J 2017;5:e3134.

Kato S, Shibuya T, Takaki Y et al. Genome-enabled metabolic reconstruction of dominant chemosynthetic colonizers in deep-sea massive sulfide deposits. Environ Microbiol 2018;20:862–77.

Kjeldsen KU, Schreiber L, Thorup CA et al. On the evolution and physiology of cable bacteria. Proc Natl Acad Sci USA 2019;116:19116–25.

Klein M, Friedrich M, Roger AJ et al. Multiple lateral transfers of dissimilatory sulfite reductase genes between major lineages of sulfate-reducing prokaryotes. J Bacteriol 2001;183: 6028–35.

Knittel K, Boetius A. Anaerobic oxidation of methane: progress with an unknown process. Annu Rev of Microbiol 2009;63:311–34.

Langwig MV, De Anda V, Dombrowski N et al. Large-scale protein level comparison of Deltaproteobacteria reveals cohesive metabolic groups. ISME J 2022;16:307–20.

Lefèvre CT, Howse PA, Schmidt ML et al. Growth of magnetotactic sulfate-reducing bacteria in oxygen concentration gradient medium. Environ Microbiol Rep 2016;8:1003–15.

Leloup J, Fossing H, Kohls K et al. Sulfate-reducing bacteria in marine sediment (Aarhus Bay, Denmark): abundance and diversity related to geochemical zonation. Environ Microbiol 2009;11:1278–91.

Löffler M, Feldhues J, Venceslau SS et al. DsrL mediates electron transfer between NADH and rDsrAB in Allochromatium vinosum. Environ Microbiol 2020;22:783–95.

Loy A, Duller S, Baranyi C et al. Reverse dissimilatory sulfite reductase as phylogenetic marker for a subgroup of sulfur-oxidizing prokaryotes. Environ Microbiol 2009;11:289–99.

Loy A, Duller S, Wagner M. Evolution and ecology of microbes dissimilating sulfur compounds: insights from siroheme sulfite reductases. In: Dahl C (ed) Microbial Sulfur Metabolism. Berlin: Springer, 2008, 46–59.

Lübbe YJ, Youn H-S, Timkovich R et al. Siro(haem)amide in Allochromatium vinosum and relevance of DsrL and DsrN, a homolog of cobyrinic acid a,c-diamide synthase, for sulphur oxidation. FEMS Microbiol Lett 2006;261:194–202.

Marietou A, Kjeldsen KU, Glombitza C et al. Response to substrate limitation by a marine sulfate-reducing bacterium. ISME J 2022;16:200–10.

McKay LJ, Dlakić M, Fields MW, et al. Co-occurring genomic capacity for anaerobic methane and dissimilatory sulfur metabolisms discovered in the Korarchaeota. Nat Microbiol 2019;4:614-22.

Medvecky M, Cejkova D, Polansky O, et al. Whole genome sequencing and function prediction of 133 gut anaerobes isolated from chicken caecum in pure cultures. BMC Genom 2018;19:561.

Mendler K, Chen H, Parks DH et al. AnnoTree: visualization and exploration of a functionally annotated microbial tree of life. Nucleic Acids Res 2019;47: 4442–8.

Momper L, Jungbluth SP, Lee MD. Energy and carbon metabolisms in a deep terrestrial subsurface fluid microbial community. ISME J 2017;11: 2319–33.

Müller AL, Kjeldsen KU, Rattei T et al. Phylogenetic and environmental diversity of DsrAB-type dissimilatory (bi)sulfite reductases. ISME J 2015;9:1152–65.

Muyzer G, Stams AJM. The ecology and biotechnology of sulphate-reducing bacteria. Nat Rev Microbiol 2008;6:441–54.

Nagakura T, Schubert F, Wagner D et al. Biological sulfate reduction in deep subseafloor sediment of guaymas basin. Front Microbiol 2022;13:845250.

Neukirchen S, Sousa FLY. DiSCo: a sequence-based type-specific predictor of Dsr-dependent dissimilatory sulphur metabolism in microbial data. Microb Genom 2021;7:000603.

Nguyen L-T, Schmidt HA, von Haeseler A et al. IQ-TREE: a fast and effective stochastic algorithm for estimating maximum-likelihood phylogenies. Mol Biol Evol 2015;32:268–74.

Oren A, Garrity GM. Valid publication of the names of forty-two phyla of prokaryotes. Int J Syst Evol Microbiol 2021;71:005056.

Parks DH, Chuvochina M, Chaumeil P-A et al. A complete domain-to-species taxonomy for Bacteria and Archaea. Nat Biotechnol 2020;38:1079–86.

Parks DH, Chuvochina M, Waite DW et al. A standardized bacterial taxonomy based on genome phylogeny substantially revises the tree of life. Nat Biotechnol 2018;36:996–1004.

Parks DH, Imelfort M, Skennerton CT et al. CheckM: assessing the quality of microbial genomes recovered from isolates, single cells, and metagenomes. Genome Res 2015;25:1043–55.

Parks DH, Rinke C, Chuvochina M, et al. Recovery of nearly 8,000 metagenome-assembled genomes substantially expands the tree of life. Nat Microbiol 2017;2:1533-42.

Pelikan C, Herbold CW, Hausmann B et al. Diversity analysis of sulfite-and sulfate-reducing microorganisms by multiplex *dsrA* and *dsrB* amplicon sequencing using new primers and mock community-optimized bioinformatics. Environ Microbiol 2016;18:2994–3009.

Pester M, Knorr K-H, Friedrich M et al. Sulfate-reducing microorganisms in wetlands - fameless actors in carbon cycling and climate change. Front Microbiol 2012;3:72.

Pereira IAC, Ramos A, Grein F et al. A comparative genomic analysis of energy metabolism in sulfate reducing bacteria and archaea. Front Microbiol 2011;2:69.

Picard A, Gartman A, Clarke DR et al. Sulfate-reducing bacteria influence the nucleation and growth of mackinawite and greigite. Geochim Cosmochim Acta 2018;220:367–84.

Pfeffer C, Larsen S, Song J et al. Filamentous bacteria transport electrons over centimetre distances. Nature 2012;491:218–21.

Probandt D, Knittel K, Tegetmeyer HE et al. Permeability shapes bacterial communities in sublittoral surface sediments. Environ Microbiol 2017;19:1584–99.

Probst AJ, Castelle CJ, Singh A et al. Genomic resolution of a cold subsurface aquifer community provides metabolic insights for novel microbes adapted to high CO_2_ concentrations. Environ Microbiol 2017;19:459–74.

Probst AJ, Ladd B, Jarett JK, et al. Differential depth distribution of microbial function and putative symbionts through sediment-hosted aquifers in the deep terrestrial subsurface. Nat Microbiol 2018;3:328-36.

Qian Z, Hao T, Mackey HR et al. Recent advances in dissimilatory sulfate reduction: from metabolic study to application. Water Res 2019;150:162–81.

Rabus R, Hansen TA, Widdel F. Dissimilatory sulfate- and sulfur-reducing prokaryotes. In: Rosenberg E, DeLong EF, Lory S et al. (eds.). The Prokaryotes: Prokaryotic Physiology and Biochemistry. Berlin, Heidelberg: Springer, 2013, 309–404.

Rabus R, Venceslau SS, Wöhlbrand L et al. A post-genomic view of the ecophysiology, catabolism and biotechnological relevance of sulphate-reducing prokaryotes. In: Poole RK (ed). Advances in Microbial Physiology. Academic Press, 2015, 55–321.

Ramel F, Brasseur G, Pieulle L et al. Growth of the obligate anaerobe *Desulfovibrio vulgaris* Hildenborough under continuous low oxygen concentration sparging: impact of the membrane-bound oxygen reductases. PLoS One 2015;10:e0123455.

Ranjan S, Todd ZR, Sutherland JD et al. Sulfidic anion concentrations on early Earth for surficial origins-of-life chemistry. Astrobiology 2018;18:1023–40.

Rey FE, Gonzalez MD, Cheng J et al. Metabolic niche of a prominent sulfate-reducing human gut bacterium. Proc Natl Acad Sci USA 2013;110:13582–7.

Rinke C, Chuvochina M, Mussig AJ, et al. A standardized archaeal taxonomy for the Genome Taxonomy Database. Nat Microbiol 2021;6:946-59.

Risgaard-Petersen N, Kristiansen M, Frederiksen RB et al. Cable bacteria in freshwater sediments. Appl Environ Microbiol 2015;81:6003–11.

Santos AA, Venceslau SS, Grein F et al. A protein trisulfide couples dissimilatory sulfate reduction to energy conservation. Science 2015;350:1541–5.

Saxton MA, Samarkin VA, Madigan MT et al. Sulfate reduction and methanogenesis in the hypersaline deep waters and sediments of a perennially ice-covered lake. Limnol Oceanogr 2021;66:1804–18.

Schoeffler M, Gaudin A, Ramel F et al. Growth of an anaerobic sulfate-reducing bacterium sustained by oxygen respiratory energy conservation after O_2_-driven experimental evolution. Environ Microbiol 2019;21:360–73.

Selma MV, Tomás-Barberán FA, Beltrán D et al. *Gordonibacter urolithinfaciens* sp. nov., a urolithin-producing bacterium isolated from the human gut. Int J Syst Evol Microbiol 2014; 64:2346–52.

Shen Y, Buick R, Canfield D. Isotopic evidence for microbial sulphate reduction in the early Archaean era. Nature 2001;410:77–81.

Sheik CS, Jain S, Dick GJ. Metabolic flexibility of enigmatic SAR324 revealed through metagenomics and metatranscriptomics. Environ Microbiol 2014;16:304–17.

Singh SB, Lin HC. Hydrogen sulfide in physiology and diseases of the digestive tract. Microorganisms 2015;3:866–89.

Stacy A, Andrade-Oliveira V, McCulloch JA et al. Infection trains the host for microbiota-enhanced resistance to pathogens. Cell 2021;184:615–27.

Stockdreher Y, Venceslau SS, Josten M et al. Cytoplasmic sulfurtransferases in the purple sulfur bacterium Allochromatium vinosum: evidence for sulfur transfer from DsrEFH to DsrC. PLoS One 2012;7:e40785.

Swan BK, Martinez-Garcia M, Preston CM et al. Potential for chemolithoautotrophy among ubiquitous bacteria lineages in the dark ocean. Science 2011;333:1296–300.

Tan S, Liu J, Fang Y et al. Insights into ecological role of a new deltaproteobacterial order Candidatus Acidulodesulfobacterales by metagenomics and metatranscriptomics. ISME J 2019;13:2044–57.

Tanabe TS, Dahl C. HMS-S-S: A tool for the identification of Sulphur metabolism-related genes and analysis of operon structures in genome and metagenome assemblies. Mol Ecol Resour 2022;22:2758–74.

Thiel J, Byrne JM, Kappler A et al. Pyrite formation from FeS and H_2_S is mediated through microbial redox activity. Proc Natl Acad Sci USA 2019;116:6897–902.

Thiel V, Garcia Costas AM, Fortney NW et al. “*Candidatus* Thermonerobacter thiotrophicus,” a non-phototrophic member of the *Bacteroidetes/Chlorobi* with dissimilatory sulfur metabolism in hot spring mat communities. Front Microbiol 2019);9:3159.

Thomsen U, Thamdrup B, Stahl DA et al. Pathways of organic carbon oxidation in a deep lacustrine sediment, Lake Michigan. Limnol Oceanogr 2004;49:2046–57.

Thorup C, Schramm A, Findlay AJ et al. Disguised as a sulfate reducer: growth of the Deltaproteobacterium Desulfurivibrio alkaliphilus by sulfide oxidation with nitrate. mBio 2017;8:e00671–17.

Urban NR, Brezonik PL, Baker LA et al. Sulfate reduction and diffusion in sediments of Little Rock Lake, Wisconsin. Limnol Oceanogr 1994;39:797–815.

van Vliet DM, von Meijenfeldt FAB, Dutilh BE et al. The bacterial sulfur cycle in expanding dysoxic and euxinic marine waters. Environ Microbiol 2021;23:2834–57.

Vigneron A, Cruaud P, Alsop E et al. Beyond the tip of the iceberg; a new view of the diversity of sulfite- and sulfate-reducing microorganisms. ISME J 2018;12:2096–9.

Wacey D, Kilburn M, Saunders M et al. Microfossils of sulphur-metabolizing cells in 3.4-billion-year-old rocks of Western Australia. Nature Geosci 2011;4:698–702.

Wagner M, Roger AJ, Flax JL et al. Phylogeny of dissimilatory sulfite reductases supports an early origin of sulfate respiration. Appl Environ Microbiol 1998;180:2975–82.

Waite DW, Chuvochina M, Pelikan C et al. Proposal to reclassify the proteobacterial classes Deltaproteobacteria and Oligoflexia, and the phylum Thermodesulfobacteria into four phyla reflecting major functional capabilities. Int J Syst Evol Microbiol 2020;70:5972–6016.

Wang Y, Wegener G, Hou J, et al. Expanding anaerobic alkane metabolism in the domain of Archaea. Nat Microbiol 2019;4:595-602.

Wasmund K, Cooper M, Schreiber L et al. Single-cell genome and group-specific dsrAB sequencing implicate marine members of the Class Dehalococcoidia (Phylum Chloroflexi) in sulfur cycling. mBio 2016;7:e00266–16.

Wasmund K, Mußmann M, Loy A. The life sulfuric: microbial ecology of sulfur cycling in marine sediments. Environ Microbiol Rep 2017;9:323–44.

Wei Y, Zhang Y. Glycyl radical enzymes and sulfonate metabolism in the microbiome. Annu Rev Biochem 2021;90:817–46.

Wolf PG, Cowley ES, Breister A et al. Diversity and distribution of sulfur metabolic genes in the human gut microbiome and their association with colorectal cancer. Microbiome 2022; 10:64.

Woodcroft BJ, Singleton CM, Boyd JA et al. Genome-centric view of carbon processing in thawing permafrost. Nature 2018;560:49–54.

Wörner S, Pester M. Microbial succession of anaerobic chitin degradation in freshwater sediments. Appl Environ Microbiol 2019a;85:e00963–19.

Wörner S, Pester M. The active sulfate-reducing microbial community in littoral sediment of oligotrophic Lake Constance. Front Microbiol 2019b;10:247.

Wu B, Liu F, Zhou A et al. Experimental evolution reveals nitrate tolerance mechanisms in Desulfovibrio vulgaris. ISME J 2020;14: 2862–76.

Würdemann D, Tindall BJ, Pukall R et al. Gordonibacter pamelaeae gen. nov., sp. nov., a new member of the Coriobacteriaceae isolated from a patient with Crohn’s disease, and reclassification of Eggerthella hongkongensis Lau et al. 2006 as Paraeggerthella hongkongensis gen. nov., comb. nov. Int J Syst Evol Microbiol 2009;59:1405–15.

Wylensek D, Hitch TCA, Riedel T et al. A collection of bacterial isolates from the pig intestine reveals functional and taxonomic diversity. Nat Commun 2020;11:6389.

Ye H, Borusak D, Eberl C et al. A novel taurine-respiring murine gut bacterium contributes to colonization resistance against enteropathogens. bioRxiv;2022. doi: https://doi.org/10.1101/2022.10.05.510937

Zecchin S, Mueller RC, Seifert J et al. Rice paddy Nitrospirae carry and express genes related to sulfate respiration: proposal of the new genus “Candidatus Sulfobium.” Appl Environ Microbiol 2018;84:e02224–17.

Zhao R, Biddle JF. Helarchaeota and co-occurring sulfate-reducing bacteria in subseafloor sediments from the Costa Rica Margin. ISME Commun 2021;1:25.

Zhou Z, Liu Y, Xu W et al. Genome-and community-level interaction insights into carbon utilization and element cycling functions of Hydrothermarchaeota in hydrothermal sediment. Msystems 2020;5:e00795–19.

Zhou Z, Tran PQ, Breister AM et al. METABOLIC: high-throughput profiling of microbial genomes for functional traits, metabolism, biogeochemistry, and community-scale functional networks. Microbiome 2022;10:33.

Zverlov V, Klein M, Lücker S et al. Lateral gene transfer of dissimilatory (bi)sulfite reductase revisited. J Bacteriol 2005;187:2203–8.

